# Poxvirus-induced concurrent activation of SAMD9 and ERK establishes a unique immune state with heightened IFN-I responses to danger signals

**DOI:** 10.64898/2026.11.22.768462

**Authors:** Tahseen Raza, Li Ma, Erich A. Peterson, Jason Liem, Jia Liu

**Affiliations:** Department of Microbiology and Immunology, University of Arkansas for Medical Sciences (UAMS), Little Rock, AR, USA; Winthrop P. Rockefeller Cancer Institute, University of Arkansas for Medical Sciences, Little Rock, AR, USA

**Keywords:** Phenylalanine, codon bias, poxvirus, SAMD9, myxoma virus, vaccinia virus, ERK1/2 pathway, type 1 interferon, Interferon-stimulated genes (ISGs)

## Abstract

Mammalian poxviruses must overcome the cellular restriction factor SAMD9 to establish productive infection. SAMD9 is a widely expressed cytoplasmic sensor of double-stranded nucleic acids (dsNA) that induces type-1 interferon (IFN-I) responses and possesses anticodon-nuclease activity targeting tRNA^phe^ specifically. To counteract SAMD9, poxviruses have evolved dedicated antagonists, including myxoma virus (MYXV) M062, which blocks both dsNA sensing and tRNA^phe^-nuclease activity. However, the significance of tRNA^phe^ depletion during poxvirus infection has remained unclear. Here, we showed that mammalian poxviruses exhibit an unexpected enrichment of phenylalanine within their proteomes and preferentially utilize UUU phenylalanine codon. In contrast, host interferon-stimulated genes (ISGs) have lower phenylalanine frequencies than poxviruses and instead preferentially utilize UUC codon. Together with the known preferentially translational impairment of UUU-rich transcripts following SAMD9-mediated tRNA^phe^ depletion, these findings suggest a mechanism by which unobstructed SAMD9 disproportionally restricts poxvirus protein synthesis while preserving host antiviral protein production. Next, using an M062-deficient MYXV mutant (Δ*M062R*), we characterized the cellular consequences of unrestricted SAMD9 activation. Unlike vaccinia virus lacking the SAMD9 antagonists C7 and K1 (VACV-Δ*C7L*/Δ*K1L*), Δ*M062R* infection induced robust ERK1/2 signaling and preserved cell growth despite SAMD9 activation. Meanwhile, Δ*M062R* infection sustained expression of antiviral and proinflammatory proteins while restricting viral protein synthesis. This established a unique cellular state characterized by heightened responsiveness to subsequent innate immune stimulation. These findings identify a previously unrecognized interaction between SAMD9-mediated translation control and ERK1/2 signaling that renders a buffering effect to catastrophic SAMD9-mediated translation shutdown and thus provide new insight into antiviral restriction and innate immune regulation.

**IMPORTANCE:** Mammalian poxviruses remain medically important to humans and veterinary hosts. Our findings provide a mechanistic explanation for the longstanding evolutionary pressure on poxviruses to antagonize SAMD9 by demonstrating a conserved dependence of poxvirus proteomes on phenylalanine-rich protein synthesis. In contrast to host interferon-stimulated genes, which contain lower frequencies of phenylalanine and preferentially utilize the UUC codon, poxviruses are enriched for phenylalanine and exhibit a strong bias towards the UUU codon. These opposing codon preferences provide a framework for understanding how SAMD9-mediated depletion of tRNA^phe^ can preferentially restrict viral protein synthesis while preserving host antiviral responses. Together these findings establish a direct mechanistic link between SAMD9-mediated antiviral restriction and innate immune regulation. Finally, identification of ERK1/2 signaling as a pathway that buffers the cellular consequences of SAMD9 activation advances our understanding of SAMD9 biology and suggests new therapeutic opportunities for diseases driven by gain-of-function SAMD9 mutations and dysregulated translation control.

## INTRODUCTION

Viruses and their hosts continuously co-evolve, resulting in diverse viral strategies that evade host restriction mechanisms. However, the immunological impact of the only known poxvirus restriction factor, SAMD9, and the cellular signaling pathways engaged following SAMD9 activation remain poorly understood.

SAMD9 is a large cytoplasmic protein that is ubiquitously expressed in immune cells and inducible upon exposure to type 1 interferon (IFN-I). SAMD9 functions as a sensor of double-stranded nucleic acids (dsNAs), including both dsDNA and dsRNA, and different viruses antagonize the SAMD9 pathway through distinct strategies. Among these viruses, mammalian poxviruses evade the SAMD9 pathway without actively degrading SAMD9 protein. Instead, poxviral inhibitors bind to SAMD9 and block its key functions. Recently, biochemical studies demonstrated the anticodon nuclease activity targeting tRNA^phe^ in the AlbA_2-like domain of SAMD9, which overlaps with the dsNA binding domain and comprises approximately 250 amino acids (residue 134-385). Remarkably, the dsNA-sensing function by SAMD9 that triggers IFN-I responses is independent from its anticodon nuclease activity.

Although the anticodon nuclease function of SAMD9 can deplete tRNA^phe^ during infection with certain poxvirus mutants lacking SAMD9 inhibitors (e.g., vaccinia virus lacking both *C7L* and *K1L*, VACV-Δ*C7L*/Δ*K1L*), this activity has not been observed in uninfected cells or in cells infected by other viruses, such as rotaviruses, which also antagonize the SAMD9 pathway. More importantly, SAMD9-mediated depletion of tRNA^phe^ exerts an unequal effect on translation of the two phenylalanine codons, UUU and UUC. Biochemical studies have shown that depletion of tRNA^phe^ by SAMD9 stalls translation of mRNAs enriched for UUU codons more severely than those enriched for UUC codons. Consequently, several important questions remain unresolved: Why have poxviruses evolved mechanisms to inhibit SAMD9 anti-codon tRNA^phe^ nuclease activity, and what are the cellular and immunological consequences of tRNA^phe^ depletion?

Given the unique and potentially profound impact of maintaining proper tRNA^phe^ levels on cellular physiology and immunity, poxviruses provide essential experimental tools for investigating these questions. Furthermore, these viral systems offer an opportunity to uncover additional cellular regulatory mechanisms to shield cells that protect cells from the potentially catastrophic consequences of translation elongation defects resulting from SAMD9-mediated depletion of tRNA^phe^.

Here, through integrated analyses of viral and host proteomes, we demonstrate that poxviruses must prevent disruption of tRNA^phe^ levels because of their elevated dependence on phenylalanine residues within the poxviral proteome. Furthermore, polarized codon usage biases, favoring UUC among host interferon stimulated genes (ISGs) and UUU among poxviral genes, suggest that SAMD9 activation may disproportionally impair viral protein synthesis while permitting continued translation of selected host protein synthesis. This observation partially explains the unusual and potent immunological state induced by myxoma virus lacking its SAMD9 inhibitor, *M062R* (Δ*M062R* MYXV), which enhances proinflammatory response upon subsequent exposure to danger signals.

However, not all poxviruses lacking SAMD9 inhibitors elicit the same cellular responses. Previous studies demonstrated that infection with VACV-Δ*C7L*/Δ*K1L* causes global inhibition of protein translation. In contrast, we found that Δ*M062R* MYXV preserves cell growth and leads to a pro-survival state, whereas VACV-Δ*C7L*/Δ*K1L* causes cell death. These observations prompted us to investigate an additional layer of cellular regulation through which SAMD9 modulates the ERK1/2 signaling pathway. We show that Δ*M062R* infection induces concurrent activation of ERK1/2 signaling and SAMD9-mediated translation repression to viral proteins, while preserving and even enhancing host synthesis of ISG-encoded proteins. This cooperative response generates a potent innate immune state that protects cells against subsequent danger signals.

## RESULTS

### Enrichment of phenylalanine in poxvirus proteomes and divergent phenylalanine codon usage between poxviruses and host ISGs

Considering the evolutionary relationship between mammalian poxviruses and SAMD9, and the ability of SAMD9 to cleave tRNA^phe^, we hypothesized that poxviruses may exhibit unique patterns of phenylalanine (Phe) utilization within their proteomes different from that of the host. To test this hypothesis, we first examined the frequency of Phe in the human proteome using UniProtKB/Swiss-Prot Release 2026_01 (28 January 2026), which contains 574,627 sequence entries comprising 208,482,574 amino acids. According to the official database statistics (Section 6.1 - Amino Acid Composition), Phe accounts for 3.86% of all amino acid residues.^1,2^

We then calculated Phe frequencies in the proteomes of myxoma virus (MYXV; accession number: AF170726.2), vaccinia virus (VACV; WR strain, AY243312.1), and monkeypox virus (MPXV; USA2022_MA_001, ON563414.3). In addition, we examined several widely recognized collections of ISGs, including those reported by Schoggins et al., 2011^3^, Schoggins et al, 2014^4^, and Yu et al., 2025^5^, to determine their Phe usage frequencies. We found that the average Phe frequency in human ISGs is lower than that observed across the human genome (**Figure 1A** and **1D**). In contrast, the proteomes of MYXV, VACV, and MPXV contain significantly higher Phe frequencies than human ISGs (**Figure 1A**). Notably, the proteome of rotaviruses (**Supplemental Table 1**) and reovirus T1L (**Supplemental Table 2**), whose dsRNA genomes are sensed by SAMD9 to induce IFN-I responses^6^, do not exhibit elevated Phe frequencies and are comparable to human ISGs (**Figure 1D**).

**Figure 1.**
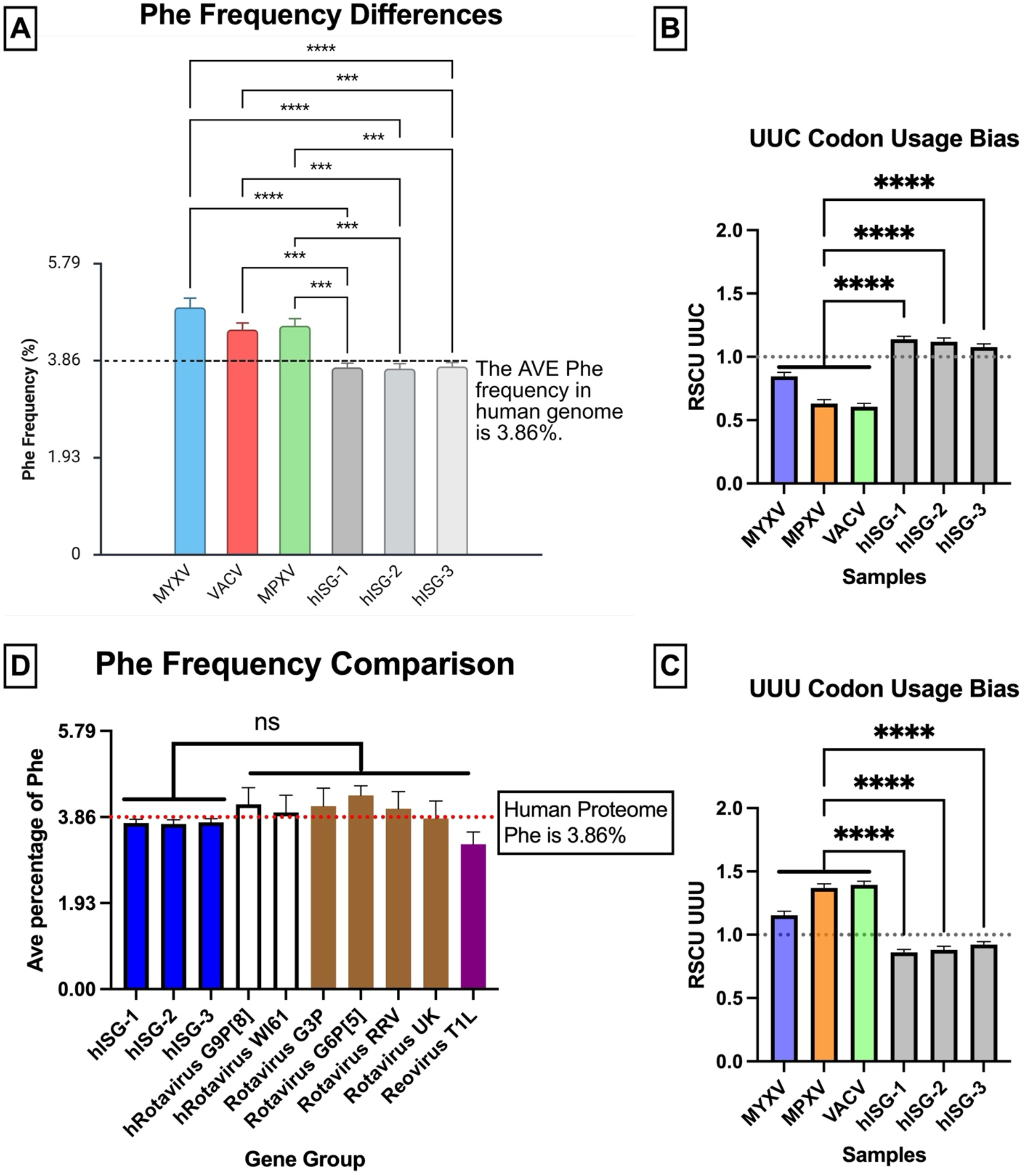
Poxvirus proteome presents phenylalanine bias and UUU codon bias for phenylalanine associated translation elongation. (A). Human ISGs show significantly lower frequency in phenylalanine usage than poxviruses. With human genome phenylalanine usage frequency of 3.86% as a baseline, representative poxviruses (MYXV, VACV, and MPXV) and human ISGs (extracted from three independent reports, Schoggins et al., 2011^3^, Schoggins et al, 2014^4^, and Yu et a., 2025^5^) diverge significantly on phenylalanine usage in corresponding proteomes. (B and C). Using RSCU to determine whether a codon usage bias is present for translation elongation to add phenylalanine into the growing polypeptide. Human ISGs prefer UUC while all representative poxviruses disfavor UUC codon usage (B). On the other hand, poxviruses show a common UUU codon bias that is not preferred by human ISGs with statistical significance (C). (D) Representative rotaviruses and reovirus^6^ do not show a clear phenylalanine preference in their proteome. Using a similar strategy as in (A), we examined viral proteome using viral genes listed in **Supplementary Table 1** and **2** and found these viral proteomes do not favor phenylalanine or significantly differ from human ISGs on phenylalanine usage.

Because the translational consequences of SAMD9 activation are determined not only by Phe abundance but also by the codons used to encode Phe, we next examined codon usage patterns. Phenylalanine is encoded by two synonymous codons: UUU and UUC. Comparative analysis of codon usage revealed a striking difference in Phe codon preference between poxviruses and human genes. All three poxviruses examined preferentially utilize UUU over UUC, whereas human ISGs display the opposite pattern, with a strong preference for UUC (**Figure 1B** and **1C**).

At first glance, such divergent codon preferences for a single amino acid appear surprising. However, closer examination of ribosome A-site occupancy under conditions of SAMD9-mediated depletion of tRNA^phe^ revealed substantially greater inhibition of translation at UUU (3.6 fold) than at UUC codons (2.6 fold).^7^ Previous work suggests that this difference arises from the biochemical properties of codon-anticodon pairing. Specifically, decoding of the UUU codon relies on a weaker non-Watson-Crick U:G interaction with Wobble-position-methylated tRNA^phe^, rendering UUU-containing transcripts more susceptible to translational repression following SAMD9 activation.^7^

These observations suggest that depletion of tRNA^phe^ following SAMD9 activation during infection with poxviruses lacking SAMD9 inhibitor(s) may impair translation of viral transcripts more severely than that of host ISG transcripts. Consequently, the opposing Phe codon usage biases of poxviruses and host ISGs may provide a mechanistic explanation for how host antiviral protein synthesis can be maintained while viral protein translation is selectively restricted during SAMD9 activation.

### Mutant ΔM062R undergoes abortive infection characterized by defective late gene expression and failure of host shutoff

We previously found that Δ*M062R* exhibits profound replication defects in virtually all mammalian cells tested, including cells derived from humans, mice, and rabbits.^8,9^ In differentiated THP-1 macrophage-like cells (**Figure 2A**), similar to observations in human HeLa cells (**Figure 2B**), Δ*M062R* likewise undergoes abortive infection. Although early viral gene expression remains largely intact,^8,10^ Δ*M062R* displays a marked defect in viral genome replication (**Figure 2C**) and fails to efficiently induce host shutoff (**Figure 2D**).

**Figure 2.**
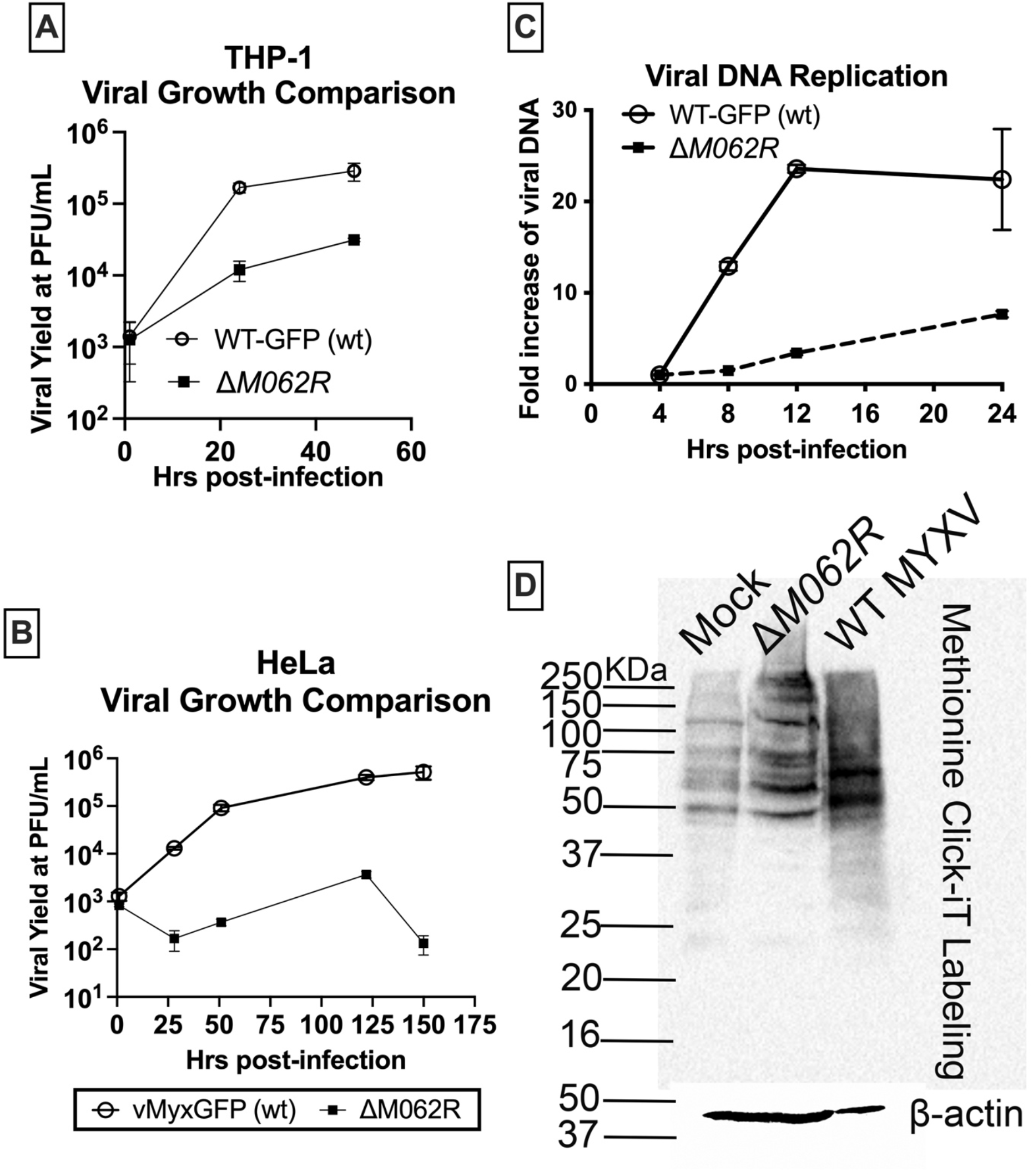
Abortive infection by D*M062R* fails to cause host shut-off with defect in viral DNA replication. (A) THP1 cells were first differentiated into macrophages before infection with either WT MYXV (WT-GFP) or Δ*M062R* at a MOI of 1. At given time points (1, 24, and 48 hrs. p.i.) cell lysates were harvested for titration on BSC-40 cells. Shown is the summary of triplicate biological replication at each time point for each virus. (B). HeLa cells were infected at a MOI of 0.2 with either WT or Δ*M062R*, and cell lysates were harvested at the given time points for titration in BSC-40 cells. (C). DNA replication defect during Δ*M062R* infection. Cells were infected with either WT or Δ*M062R* MYXV at a MOI of 5. At the given time points, DNA extraction was performed followed by real-time PCR. Relative real-time PCR was performed and fold induction calculation was conducted using ΔΔCt method as previously described^8^. (D). Infection by Δ*M062R* cannot cause host shut-off with intact host gene translation initiation. THP1-differentiated macrophages were mock treated, infected with Δ*M062R*, or WT MYXV at a MOI of 5. At 16 hrs. p.i., cells were pulse-labelled for de novo synthesized polypeptide with AHA using Click-iT chemistry. Total proteins from each group were then separated on SDS-PAGE for Western blot. Cells from Δ*M062R* infection showed the same banding pattern as mock infected cells, while WT MYXV infection caused distinct pattern for protein mobility suggesting the successful shutoff of host protein synthesis and transition to predominantly viral protein synthesis. Loading of comparable quantity of total protein is indicated by β-actin levels. Shown is the representative of three independent experiments.

To further characterize the molecular basis of this replication defect, we performed Illumina RNA sequencing to examine viral and host transcript levels during a time course of infection in differentiated THP-1 cells infected with Δ*M062R*. Using WT MYXV infection as a control, we found that Δ*M062R* infection did not cause a detectable reduction in intermediate viral gene transcription (**Supplemental Figure S1**).

We next investigated whether intermediate protein synthesis was affected during Δ*M062R* infection. To measure intermediate protein synthesis, we utilized a well-established luciferase reporter assay in which luciferase expression was driven by characterized VACV intermediate promoters, including the *I1L* promoter (p*I1L*)^11^ (**Figure 3A**) and the *A2L* promoter (p*A2L*)^11^ (**Figure 3B**). Reporter plasmids were transfected 16-24 hours (hrs.) before infection with either WT MYXV or Δ*M062R* at an MOI of 5 in the presence or absence of AraC, an inhibitor of DNA replication.

**Figure 3.**
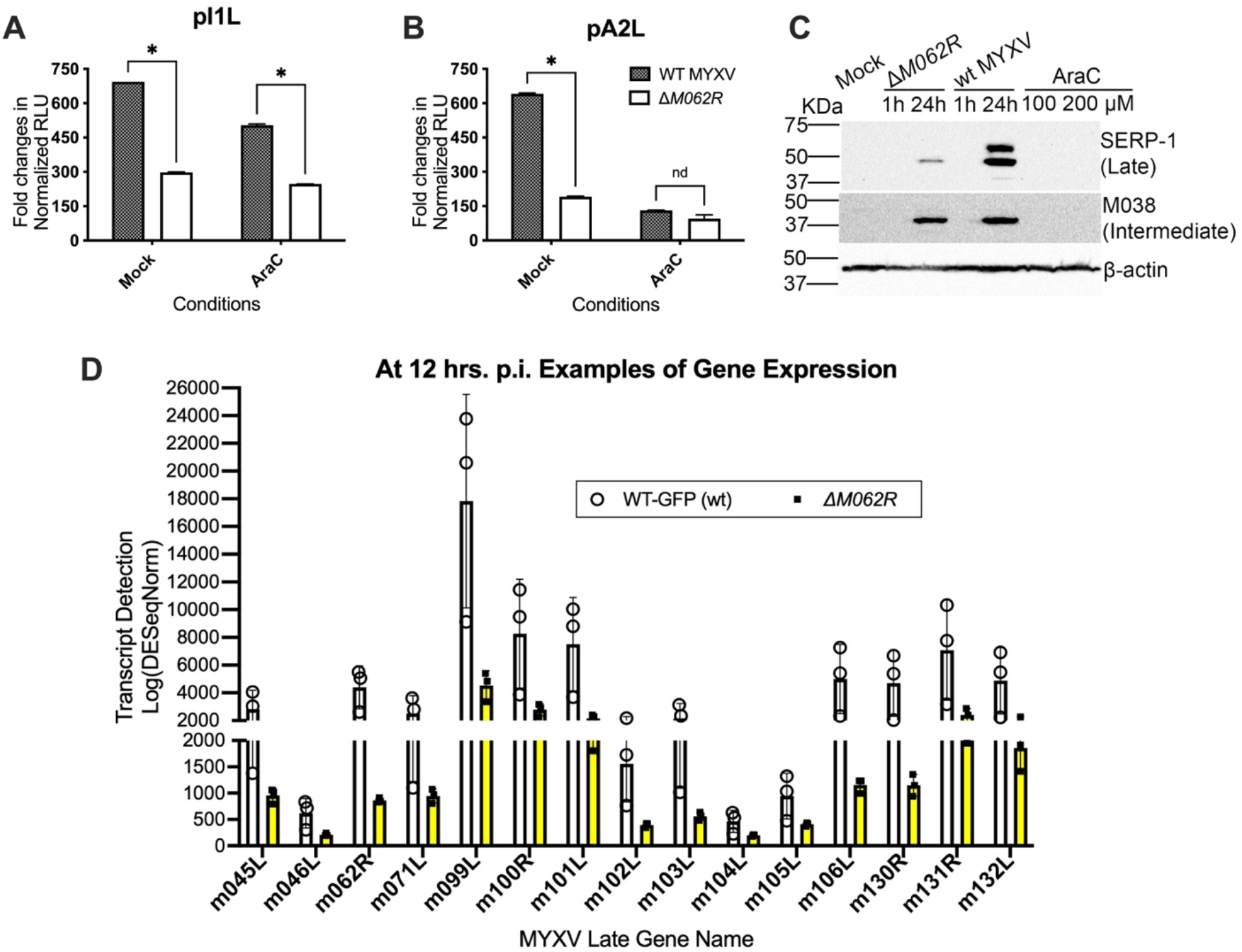
Infection by Δ*M062R* does not cause noticeable defect in intermediate protein synthesis but leads to apparent deficit in late protein synthesis due to reduced late RNA synthesis. After infection of cells with either WT MYXV or Δ*M062R* at a MOI of 5, in the presence or absence of DNA replication inhibitor, AraC (200 μM), transfection of viral intermediate promoter driven luciferase plasmid was performed using ViaFect, approximately 18 hrs. post transfection, supernatant was collected for luciferase activity assay. Results for pI1L (A), pA2L (B), and pINT (C) -driven luciferase activities are shown. Consistent results were observed using HeLa and HEK293T cells. Shown is a representative result using HeLa cells. (D) THP1 were differentiated into macrophages before mock treated, infected with Δ*M062R* or wildtype MYXV. At the given time points of 1 and 24 hrs. p.i., viral post-replicative gene expression was examined. The expression of either M038 (intermediate gene) or SERP-1 (late gene) was assessed in Western Blot. In the presence of AraC, post-replicative protein synthesis was effectively inhibited, which confirmed the kinetic group of M038 and SERP-1. (E). Late gene transcription in Δ*M062R* infection is affected relative to WT viral infection. Differential expression between WT MYXV and Δ*M062R* (n=3 biological replicate per group) was used to assess DESeq2 with raw gene-level count matrices normalized to correct for library size and compositional biases. Genes with an adjusted P-value<0.05 were considered significantly differentially expressed.

Comparison of luciferase activities between WT MYXV and Δ*M062R* infections provided an overall measure of intermediate protein synthesis during mutant virus infection. Furthermore, comparison of results obtained in the presence and absence of AraC allowed us to evaluate the contribution of viral genome replication defects to intermediate protein synthesis. Notably, inhibition of DNA replication by AraC reduced the differences in intermediate protein expression observed between WT and Δ*M062R* infections (**Figure 3A-B**). Overall, we observed a modest reduction in intermediate protein synthesis during Δ*M062R* infection relative to WT infection (**Figure 3A-B**). Consistent with the reporter assays, the modest reduction in intermediate protein synthesis during Δ*M062R* infection was not readily detectable by western blot analysis (**Figure 3C**), as illustrated by comparable expression of the intermediate viral protein M038. In contrast, both viral late-gene transcription (**Figure 3D**) and late protein production (**Figure 3C**, SERP1) were substantially impaired during Δ*M062R* infection. These observations are consistent with our previous findings demonstrating that Δ*M062R* is defective in late stages of the viral replication cycle.

SAMD9 is currently the only host factor known to mediate the abortive-infection phenotype of Δ*M062R* in human cells, as deletion of SAMD9 alone fully restores Δ*M062R* replication to levels comparable to those of WT virus.^9,12^ Collectively, these findings are consistent with a model in which unrestricted SAMD9 engages viral dsDNA during Δ*M062R* infection, resulting in activation of its anticodon nuclease activity and subsequent depletion of tRNA^phe^.^7^ Reduced availability of tRNA^phe^ would be expected to disproportionately impair translation of Phe-rich viral proteins, thereby slowing intermediate protein synthesis, restricting late gene expression, preventing efficient host shutoff, and ultimately leading to abortive infection.

### Infection by ΔM062R preserves host protein synthesis and supports expression of antiviral and proinflammatory proteins

A previous study using VACV lacking the SAMD9 inhibitors *C7L* and *K1L* (VACV-Δ*C7L/*Δ*K1L*) demonstrated that infection by this mutant causes global translational inhibition in non-immune cells such as HeLa.^7,13^ This translational shutdown^13^ is mediated by depletion of tRNA^phe^ following SAMD9 activation.^7^ In our previous work, we showed that VACV-Δ*C7L/*Δ*K1L* activates SAMD9 in a manner similarly to that observed during MYXV Δ*M062R* infection, leading to the formation of antiviral granules.^9^ Furthermore, we demonstrated that M062 protein directly binds to the dsNA-binding AlbA_2-like domain of SAMD9^12^, which overlaps with the domain responsible for the anticodon nuclease activity targeting tRNA^phe^.^7^

Based on these observations, it is reasonable to hypothesize that infection with Δ*M062R* would likewise result in widespread suppression of host protein synthesis. However, Δ*M062R* infection also induces robust IFN-I responses and ISG expression in monocytes/macrophages^10^, a phenomenon that appears inconsistent with a model of global translation suppression. To better understand the contribution of SAMD9 tRNA^phe^ nuclease activity to innate immune responses, we examined host protein expression at 18-24 h post-infection with Δ*M062R*. Instead, we observed no detectable reduction in the abundance of any host proteins examined relative to uninfected cells (**Figure 4**). Similar findings were obtained for antiviral proteins and inflammatory cytokines in human monocytic THP1 differentiated macrophages (**Figure 4A**), primary human CD14^+^ monocytes (**Figure 4B**), and primary human dermal fibroblasts (NHDFs) (**Figure 4C**).

**Figure 4.**
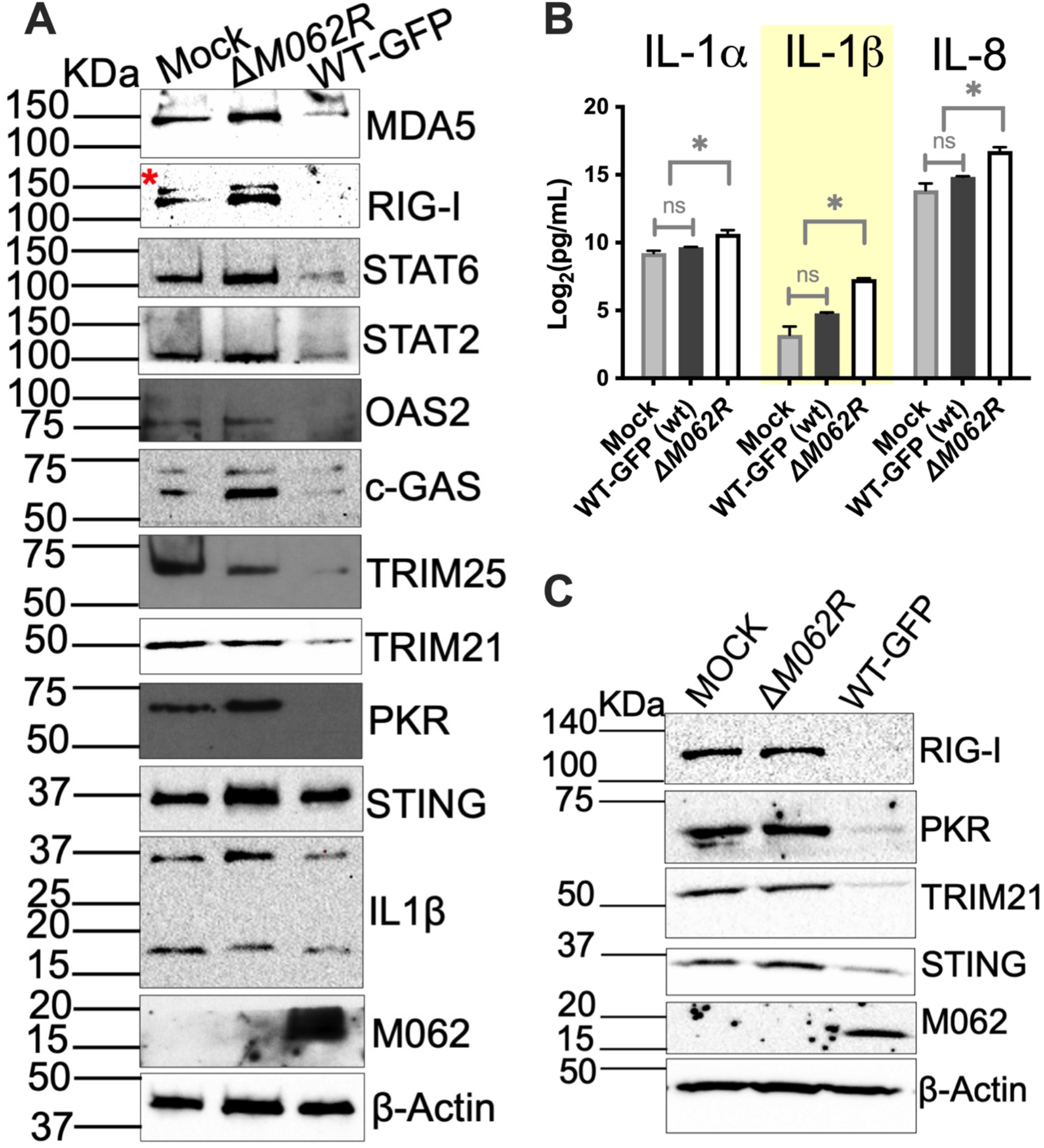
Infection by Δ*M062R* leads to continuous accumulation of host proteins including ISGs. (A). Intact host protein synthesis in differentiated human THP1 cells after Δ*M062R* treatment. Differentiated THP1 cells were mock treated, infected with Δ*M062R*, or WT MYXV. At 24 hrs. post-treatment, numbers of host protein levels were compared among three groups. A subset of antiviral proteins is specifically upregulated relative to WT MYXV infection. *The red asterisk marks the faint band above the dominant band of RIG-I in each lane; this was caused by the prior probing of MDA-5 (135 KDa) that was incompletely stripped before probing for RIG-I (102 KDa). (B) Detection of secreted proinflammatory IL1b from primary human macrophages. In primary healthy human CD14^+^ monocytes, Δ*M062R* treatment also led to upregulation of IL1a, IL1β, and IL8 detected by multi-plex immunoassay, which was consistent with what was seen in protein expression profile examined in differentiated THP1 cells in “(A)”. (C) Uninterrupted host defense molecular expression is detected in Δ*M062R* treated primary NHDF cells. At 24 hrs. post-treatment, immunoblotting was performed. Infection by WT MYXV led to depletion of host proteins tested, while Δ*M062R* did not hinder these protein syntheses.

The proteins examined consisted primarily of antiviral and pro-inflammatory proteins, many of which are established ISGs. In addition, expression of housekeeping genes such as β-actin (**Figure 4**) also remained unaffected. Thus, despite activation of SAMD9 antiviral functions and the likely induction of anticodon nuclease activity targeting tRNA^phe^, Δ*M062R*-infected cells exhibited no measurable impairment in the synthesis of the host proteins examined.

Notably, Δ*M062R* infection appeared to increase expression of a subset of host proteins, particularly antiviral and proinflammatory effectors. These findings are consistent with our earlier prediction that host ISGs should be less sensitive to fluctuations in tRNA^phe^ availability than poxviral proteins because of their distinct Phe codon usage patterns. Together, these findings suggest that SAMD9 activation establishes a selective translational environment, in which viral protein production is restricted while preserving, and in some cases enhancing, expression of host antiviral and proinflammatory genes.

### Infection by ΔM062R induces a pro-survival and growth-associated transcriptional program linked to ERK1/2 activation

We previously reported that the transcriptional profile induced by Δ*M062R* differs substantially from that elicited by canonical dsDNA-triggered IFN-I responses^10^. However, the biological significance of these differences has not been fully characterized. To further investigate the host response to Δ*M062R* infection, we analyzed the host transcriptomic profile at later time points during infection (**Figure 5A**). This analysis revealed coordinated upregulation of genes associated with cell growth and survival (**Figure 5B**).

**Figure 5.**
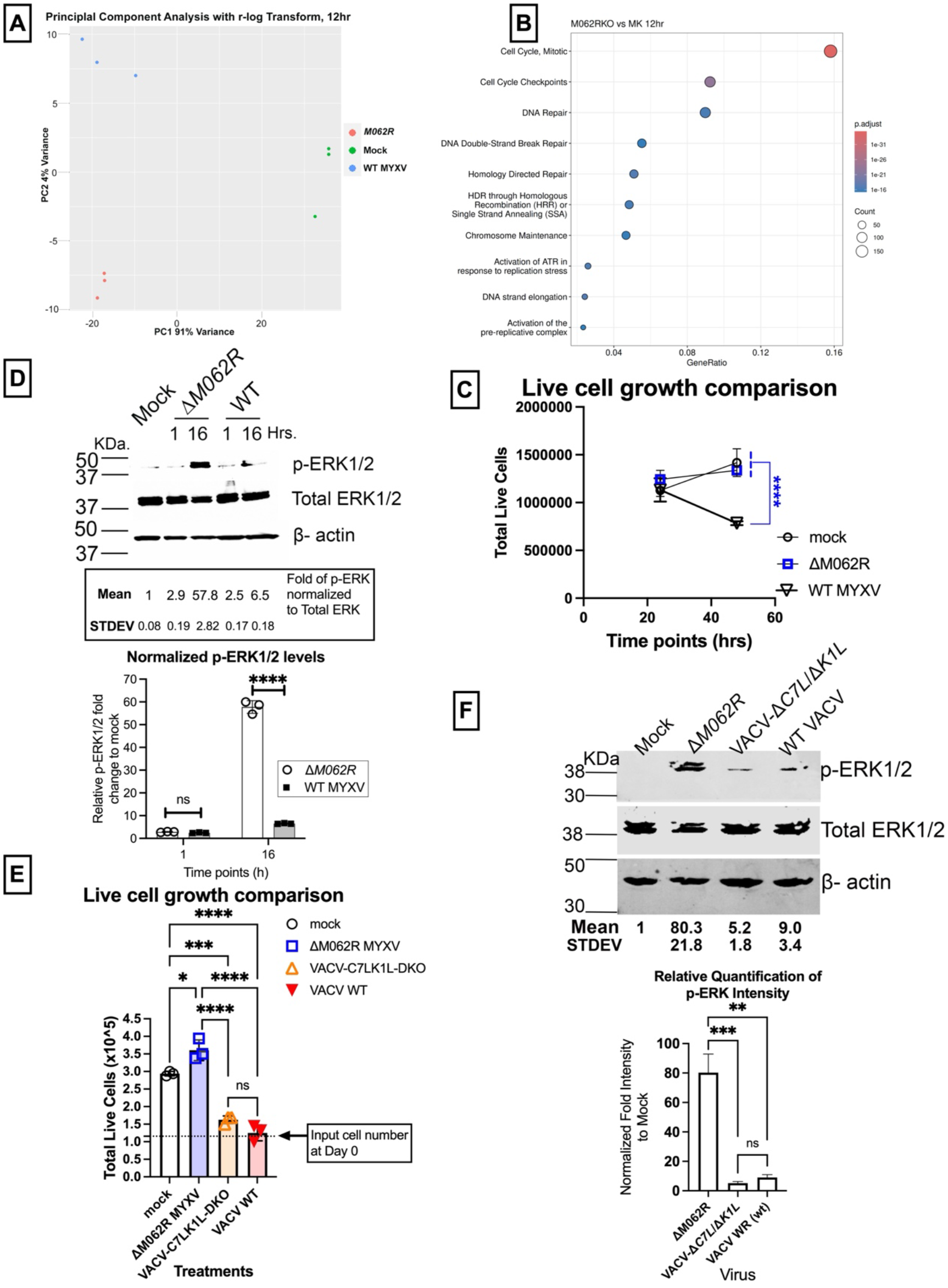
Treatment with Δ*M062R* leads to pro-survival and -growth transcriptional profile consistent with activation of ERK signaling and cell growth observed in treated cells. (A) PCA plot of RNAseq data showing well clustered treatment groups of mock treated, WT, and Δ*M062R* infection. Data from three groups of samples at 12 hours post-treatment are well separated from one another, and data within each group are clustered together. (B) Treatment by Δ*M062R* led to a unique transcriptional profile. At 12 hrs. p.i., relative to WT MYXV treatment, Δ*M062R* induced gene expression determined by GO biological processes enrichment analysis of differentially expressed genes from RNAseq data analyses. A heatmap displays the z-score standardized expression of the genes within a given term, blue for lower normalized expression and red for higher. (C). Treatment with Δ*M062R* does not hinder cell proliferation. THP1 cells that are mock treated, treated with WT or Δ*M062R* at a MOI of 1 are followed up on their growth at given time points using trypan blue exclusion test. Live cells at given times are recorded to build the growth comparison with triplicate technical replicates. Shown is the representative of 3 biological replicates. (D). Treatment of Δ*M062R* induces phosphorylation of ERK1/2 signaling. THP1 differentiated macrophages are mock treated or infected with WT or Δ*M062R* at a MOI of 5-10, and cell lysates are harvested for immunoblotting. Equal amount of total protein (50 mg) per sample is separated on SDS-PAGE for immunoblotting and probed for phosphorylated ERK1/2, total ERK1/2, and b-actin. Pixel density of detected proteins is quantitated using ImageStudio and quantitation of phosphorylated ERK is normalized to total ERK for quantitative comparison. Ratio in mock infected cells is standardized as 1 and ratio for infected cells at various time points are compared to mock infected cells for fold changes. Standardized ratio and standard deviation are calculated from triplicate technical replications. Results shown is the representative immunoblotting of 3 biological replicates. Multiple paired t-test was conducted, and statistical significance is defined as *p<0.05, ***p<0.001. (E). In contrast to Δ*M062R* promoted cell growth, treatment of VACV-D*C7L*/D*K1L*-DKO does not promote cell growth. A similar growth comparison experiment as described in “(C)” is conducted using THP1 cells with mock infection, treatment of Δ*M062R*, VACV-D*C7L*/D*K1L*-DKO, or WT VACV. One-way ANOVA followed by multiple comparisons is conducted to determine statistical significance in differences on cell growth, *p<0.05, ***p<0.001, ****p<0.0001, and ns: not significant. (F). In contrast to Δ*M062R* induced activation of p-ERK1/2 signaling, VACV-D*C7L*/D*K1L* infection does not cause detectably elevated p-ERK1/2 signaling. THP1-differentiated macrophages are mock treated or treated with Δ*M062R*, VACV-D*C7L*/D*K1L*, or WT VACV at a MOI of 5. Cell lysates are harvested for protein extraction at 9 hours post-treatment, and an equal amount of total protein (50 mg) is separated for SDS-PAGE followed by immunoblotting. Quantitation of protein detection and normalized comparison are conducted similarly as described in “(D)”. One-way ANOVA followed by multiple-comparisons is conducted to compare fold changes in p-ERK1/2 signaling, *p<0.05, **p<0.01, and ***p<0.001.

To determine whether this transcriptional signature has functional consequences, we examined the effect of Δ*M062R* infection on cell growth in human monocytic THP-1 cells. Remarkably, cells infected with Δ*M062R* exhibited growth kinetics comparable to those of uninfected cells, whereas WT MYXV infection resulted in the expected decline in cell growth and viability (**Figure 5C**).

To identify signaling pathways that might underlie this phenotype, we performed an initial targeted signaling screen, which identified ERK1/2 as a candidate pathway for further investigator. We subsequently confirmed that Δ*M062R* infection induces robust ERK1/2 activation in monocytes/macrophages (**Figure 5D**). Similar ERK1/2 activation was observed in additional cell types, including HeLa (**Supplemental Figure S2**).

We previously reported that both Δ*M062R* and VACV-Δ*C7L/*Δ*K1L* activate SAMD9-mediated antiviral responses^9^. Furthermore, VACV-Δ*C7L/*Δ*K1L* infection has been shown to induce tRNA^phe^depletion^7^ resulting in atypical global translation inhibition^13^. Indeed, infection with VACV-Δ*C7L/*Δ*K1L* inhibited cell growth to a degree comparable to that observed during WT VACV infection, in stark contrast to the growth-preserving phenotype induced by Δ*M062R* (**Figure 5E**). Moreover, VACV-Δ*C7L/*Δ*K1L* failed to induce the robust ERK1/2 activation observed during Δ*M062R* infection (**Figure 5F**).

Together, these findings suggest that activation of SAMD9 and the resulting effects on translation are not the sole determinants of the distinct cellular outcomes induced by these two mutant viruses. Although both Δ*M062R* and VACV-Δ*C7L/*Δ*K1L* lack viral SAMD9 inhibitors and therefore permit activation of SAMD9-associated antiviral functions, Δ*M062R* uniquely triggers robust ERK1/2 activation together with a pro-survival and growth-associated transcriptional program. These observations indicate that additional regulatory mechanisms beyond SAMD9-mediated tRNA^phe^ depletion contribute substantially to determining cellular fate following infection.

### Treatment of ΔM062R establishes a potentiated antiviral state associated with ERK signaling

Because ERK1/2 signaling is a well-established regulator of cell survival, proliferation, and protein synthesis, we investigated whether ERK1/2 activation contributes to the cellular responses induced by Δ*M062R* infection. In addition to its cytoplasmic functions, the ERK1/2 pathway also regulates nuclear events through the translocation of phosphorylated ERK into the nucleus, where it activates transcription factors such as ELK1^14^, which emerged as a predicted upstream regulator in the bioinformatic analysis of the transcriptomic data. Pharmacological inhibition of ERK signaling using the MEK1/2 inhibitor U0126 during Δ*M062R* infection resulted in loss of activation of multiple well-established downstream effectors. These downstream effectors of ERK1/2 involved in protein synthesis and cellular growth (**Figure 6A**), including p90RSK^15,16^, the MNK1/eIF4E axis^16–18^, and ELK-1 that controls transcriptional programs to regulate cell survival and proliferation^19,20^.

**Figure 6.**
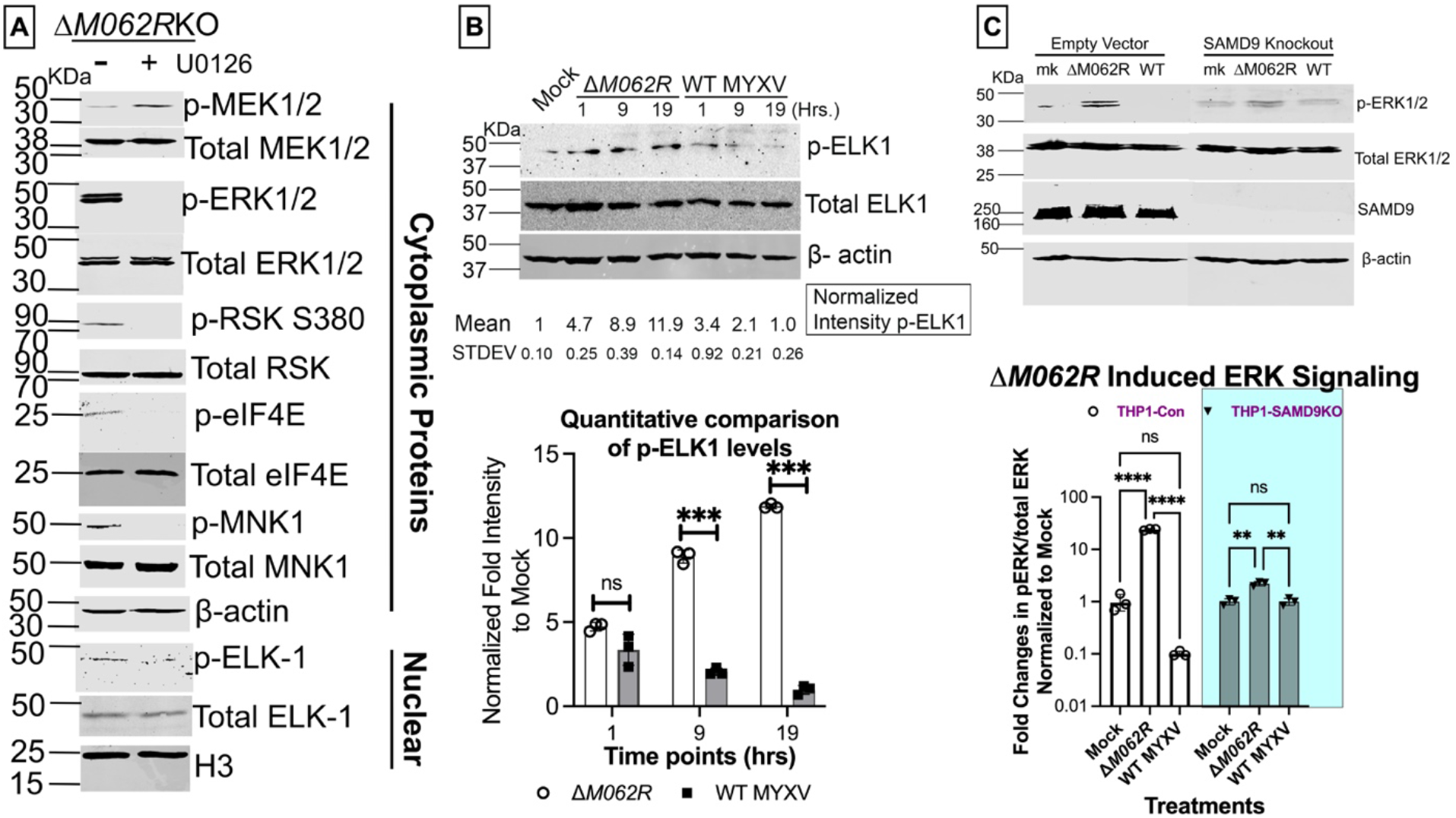
Treatment of D*M062R* activates ERK signaling that turns on effectors in both cytoplasm and nucleus. (A) Treatment with the inhibitor of ERK1/2 signaling relieves Δ*M062R*-associated activation of downstream effectors in both cytoplasm and nucleus. An inhibitor of upstream kinase of ERK1/2, U0126, was absent or present for 3 hours in Δ*M062R* treated THP1 macrophages (15 hrs. p.i.) before fractionation to isolate cytoplasmic and nuclear proteins for immunoblotting. Equal amounts of total protein (30 mg for cytoplasmic protein and 15 mg for nuclear protein) are loaded for protein separation on SDS-PAGE for immunoblotting of phosphorylated and total protein of ERK substrates. Both b-actin and H3 serve as internal loading control for cytoplasmic and nuclear proteins, respectively. (B). Treatment of Δ*M062R* leads to significant activation of ELK-1 transcription factor. Whole cell lysate of THP1 macrophages with mock treated, infection at an MOI of 5 by Δ*M062R* or WT MYXV at 1, 9, and 19 hrs. p.i. are harvested for immunoblotting. Phosphorylated and total ELK-1 along with internal loading control (b-actin) are probed for later quantitation as described above. Multiple paired t tests are performed to compared fold changes of normalized ratio of p-ELK-1/total ELK-1 to mock infected cells, ns: not significant, ***p<0.001. (C). In SAMD9-null cells, Δ*M062R* induced ERK1/2 signaling is not significantly different from the controls. In lentivirus transduced control or SAMD9-null cells, cell lysates with mock infection and infection with either Δ*M062R* or WT MYXV are separated for immunoblotting to probe for phosphorylated and total ERK1/2 along with the internal loading control (b-actin). All the data shown is a representative of 3 biological replicates.

Consistent with the activation of ERK downstream effectors during Δ*M062R* infection, ELK-1 phosphorylation was markedly increased relative to both mock-infected cells and cells infected with WT MYXV (**Figure 6B**). This observation independently validated the transcriptomic prediction that ELK-1 is a shared regulatory factor among genes upregulated during Δ*M062R* infection. These observations suggest that Δ*M062R*-induced ERK signaling engages a broad network of downstream effectors that collectively establish a distinct cellular environment characterized by unique transcriptional and translational programs.

To determine whether ERK activation was linked to SAMD9 function, we examined ERK signaling in previously generated control and SAMD9-null THP1 cells^6^. Loss of SAMD9 expression abolished the differential ERK activation during Δ*M062R* infection relative to controls (**Figure 6C**). These findings indicate that ERK activation during Δ*M062R* infection is dependent on SAMD9.

If Δ*M062R* infection promotes expression of antiviral sensors and effectors while simultaneously maintaining cellular growth and homeostasis, such a state would be predicted to enhance host defense capacity. To investigate the immunological consequences of this phenotype, we evaluated how Δ*M062R* pre-treatment affects cellular responsiveness to a subsequent innate immune stimulus.

Using a previously described dual-luciferase reporter system in THP-1^21^ as a surrogate measure of IRF-dependent IFN-I induction^10^, cells were first infected with Δ*M062R*, WT MYXV, or mock treated and subsequently challenged with either reovirus TD3 (**Figure 7A**), which activates both RIG-I and MDA5^22^, or 5’-ppp-dsRNA, a synthetic RIG-I agonist (**Figure 7B**). As expected, WT MYXV pre-treatment effectively suppressed IFN-I induction in either case. In contrast, cells pretreated with Δ*M062R* consistently displayed enhanced reporter activity compared with both mock-treated and WT MYXV-infected controls.

**Figure 7.**
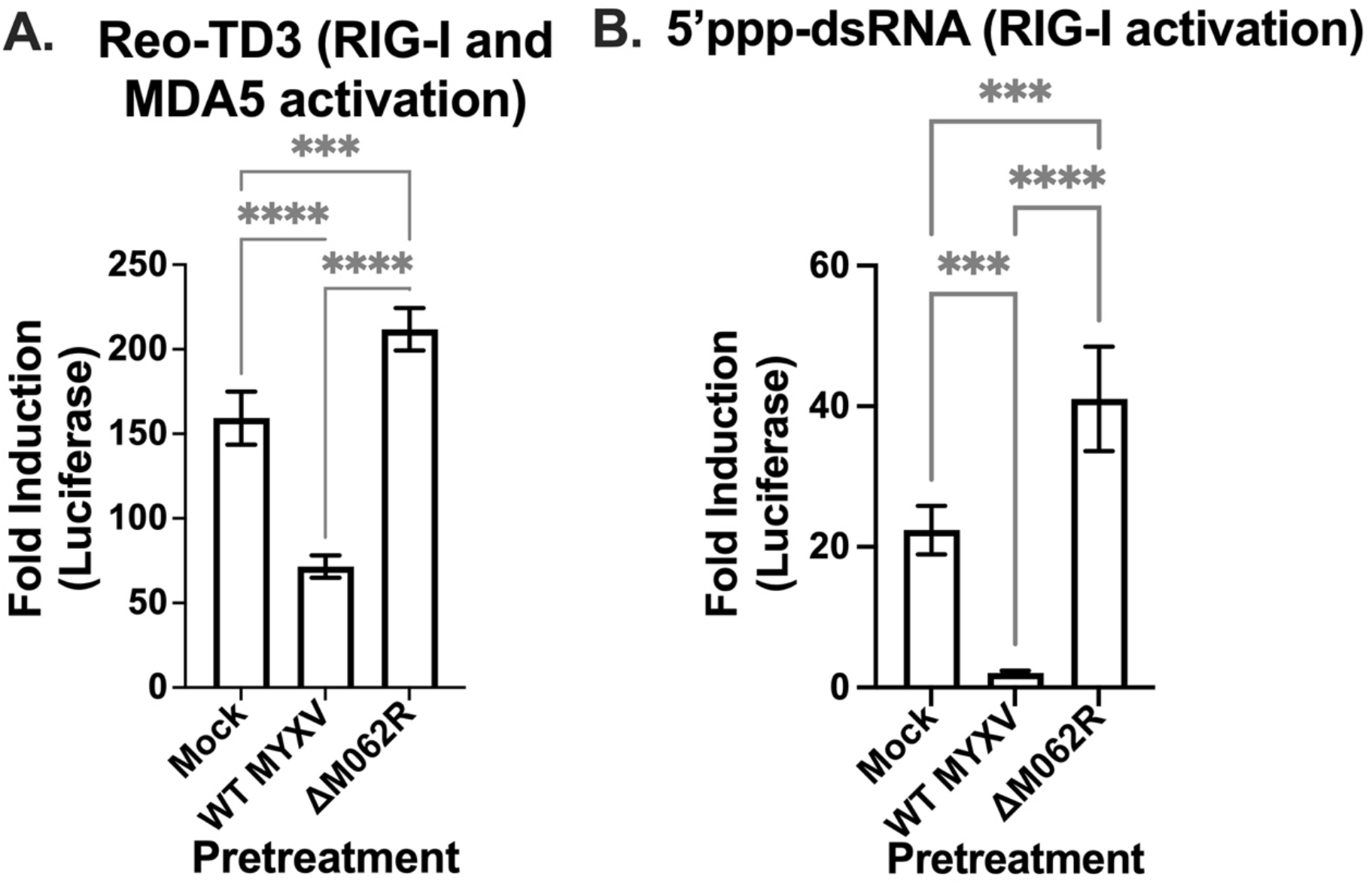
Pretreatment of Δ*M062R* potentiates macrophages for heightened IRF-dependent response upon new exposure to a danger signal. (A) Pretreatment of macrophages with Δ*M062R* leads to significantly elevated IFN-I responses upon subsequent Reo-TD3 exposure. THP1 macrophages with IRF-driven luciferase is pretreated with mock infection, infection with WT or Δ*M062R* MYXV at an MOI of 5 for 9 hrs before exposure to Reo-TD3 at an MOI of 50 for 12 hrs before measuring luciferase activities in the supernatants. Reo-TD3 is a known stimulus to both RIG-I and MDA5 leading to IFN-I induction. Shown is the summary of 2 biological replicates. (B). Pretreatment with Δ*M062R* promotes significantly higher IFN-I responses upon 3’ppp-dsRNA stimulation. Similar to experimental timeline shown in “(A)” with exception of the subsequent exposure to 3’ppp-dsRNA by transfection, at 12 hrs. post 3’ppp-dsRNA transfection, luciferase activity is measured for fold induction. Statistical comparison for both experiments is conducted using one-way ANOVA followed by multiple comparisons, ***p<0.001, and ****p<0.0001.

These findings demonstrate that Δ*M062R* infection establishes a cellular state that is primed for heightened antiviral responsiveness upon subsequent exposure to innate immune stimuli. Together with the observed activation of SAMD9-dependent ERK signaling, these results support a model in which Δ*M062R* infection induces a unique immune-potentiating state characterized by preserved cellular homeostasis, sustained expression of antiviral protein, and heightened responsiveness to subsequent danger signals.

## DISCUSSION

Viral genomes frequently exhibit distinctive nucleotide compositions that shape codon usage and, consequently, influence amino acid composition within viral proteome. For instance, both human immunodeficiency viruses (HIVs) possess strong adenine-rich nucleotide biases compared with relatively neutral composition of human T-cell leukemia viruses (HLTVs), which significantly distort viral amino acid usage to ones with codons more frequently with adenines.^23^ Viral mutation spectrum along with host selection forces (e.g., ZAP-mediated CpG suppression^24,25^, and APOBEC3s induced C->T/G->A mutagenesis^26,27^), further drives amino acid composition^28,29^, especially among RNA viruses, such as SARS-CoV2^30^. While pathogen genomes are often shaped by a persistent directional mutational pressure^31,32^ leading to equilibrium with AT-rich genome that generally skews towards codon usage of lysine, asparagine, isoleucine, tyrosine, and phenylalanine.^33–35^ Several pathogens have known amino acid bias in their proteome, e.g., asparagine in plasmodium falciparum^36–38^ and bias of isoleucine and lysine in Candidatus Zinderia insecticola^39^. However, the biological significance of virus-specific amino acid bias has received comparatively less attention. Poxviruses are large DNA viruses with their life cycle exclusively in the cytoplasm and possess unusual AT-rich genomes in majority of sequenced poxviruses^40,41^. Although it is predicted to present distinct bias towards a group of codon usage between AT- and GC-rich genomes, it was not known whether amino acid bias is present for these viruses. Previous study shows that MYXV has less AT-content in its genome than most poxviruses^41^. Thus, the finding of common bias on phenylalanine usage by MYXV comparable to VACV and MPXV is intriguing. The observation that MYXV, VACV, and MPXV all exhibit elevated phenylalanine content despite differences in overall genome composition suggests that phenylalanine enrichment may represent a conserved and biologically relevant feature of mammalian poxviruses rather than a simple consequence of genomic AT-rich content.

Viruses that encode SAMD9 antagonists^8,6^ provide valuable tools for understanding SAMD9 biology. While SAMD9 actively triggers antiviral IFN-I effect against viruses with dsRNA genomes such as rotavirus and reovirus, only rotaviruses possess inhibitors of SAMD9. More importantly, these viruses are not vulnerable to SAMD9’s tRNA^phe^ nuclease function^6^. This is consistent with the finding that both reoviruses and rotaviruses do not exhibit an apparent preference in phenylalanine usage in their proteome. On the other hand, poxviruses encode many inhibitors of IFN-I responses while also encoding multiple SAMD9 antagonists, among which so far one inhibitor—MYXV M062^8,12,10^—directly blocks the SAMD9 domain binding to dsNA that overlaps to the region possessing tRNA^phe^ nuclease activity. Other poxviral antagonizers of SAMD9, such as VACV C7 and K1, also interfere with SAMD9 tRNA^phe^ nuclease-associated translation control^7,42,43^. Together with our finding that poxvirus proteomes are enriched for phenylalanine, these results provide a potential explanation for the strong selective pressure on poxviruses to antagonize SAMD9. These observations are consistent with the possibility that selective pressures associated with SAMD9 have contributed to the maintenance of elevated phenylalanine usage among most mammalian poxviruses.

A second key finding of this study is that SAMD9-mediated depletion of tRNA^phe^ does not necessarily cause global translation collapse. In theory, activating SAMD9 tRNA^phe^ nuclease stalls translation at the elongation step, as others showed that in infected cells between 2-8 hours post-infection VACV-Δ*C7L/*Δ*K1L* caused ∼50% reduction in tRNA^phe^ levels^7^. It is thus plausible that continuous tRNA^phe^ depletion under this circumstance leads to catastrophic cell death. Our surprising results on MYXV Δ*M062R*-induced differential impact on host and viral protein synthesis expand our understanding on SAMD9-associated translation control and prompt us to reconsider the early assessment. Our findings suggest that SAMD9-mediated depletion of tRNA^phe^ is unlikely to be the sole determinant of cellular fate following infection. Instead, cellular responses appear to be shaped by additional signaling pathways capable of buffering the consequences of translation stress. In this study, we identified the ERK1/2 signaling pathway as a candidate mechanism that counterbalances translational stress associated with SAMD9 activation. For instance, the integrated stress response (ISR) is specifically induced by VACV-Δ*C7L/*Δ*K1L*^7^. With a counterbalance regulation, cell fate is not singularly determined by tRNA^phe^ availability. We acknowledge the limitation of this study that we cannot determine whether tRNA^phe^ depletion due to SAMD9 activation induced by mutant poxviruses lacking SAMD9 inhibitors is temporary or continuous. This determination is key to reveal the relationship between the SAMD9 pathway (e.g., SAMD9 conformation change-associated self-inhibition) and the cellular counterbalance with a buffering effect to maintain cellular homeostasis, and more importantly, how tRNA^phe^ nuclease activity and dsNA sensing-induced antiviral responses are coordinately regulated. Future studies will be necessary to define how SAMD9 activation, tRNA^phe^ depletion, translational stress responses, and ERK signaling are coordinated over time.

Our results suggest that the contrasting effects of Δ*M062R* and VACV-Δ*C7L/*Δ*K1L* on cell growth may, at least in part, reflect their distinct effects on the ERK signaling axis despite activating the same SAMD9 pathway. VACV encodes many viral proteins that effectively inhibit ERK signaling in human cells due to the presence, such as *M2L* as a known direct ERK inhibitor^44,45^, *A46R* that simultaneously blocks several kinases including ERK^46^, and *A52R*^47^ that suppresses ERK signaling via inhibition of TRAF6-TAK1-ERK axis^48^. On the other hand, MYXV does not encode the homolog of VACV *A46R*^49^; although MYXV possesses some putative homologues of VACV ortholog proteins, e.g. *M154L* as VACV *M2L* homolog (50% identity) and *M136R* as VACV *A52R* homolog (23% identity), they likely do not function effectively against human cellular targets^50^. These virus-specific differences may contribute to the divergent cellular responses elicited by VACV-Δ*C7L/*Δ*K1L* and Δ*M062R*. Importantly, our findings suggest that concurrent ERK1/2 activation pathway may provide a buffering mechanism to mitigate the translational consequences of SAMD9 activation.

Activation of ERK1/2 initiates signaling events in both cytoplasm and nucleus. These events promote cell growth and counteract cell stress response. We found previously that Δ*M062R* does not trigger phosphorylation of eIF2α^9^, but infection by VACV-Δ*C7L/*Δ*K1L* leads to stress response associated with activation of eIF2α (data not shown and^7^), which is consistent with our finding in this study. In addition to supporting host protein synthesis during Δ*M062R* infection, phosphorylated ERK translocates to the nucleus, where it activates transcription factors such as ELK-1. Motif enrichment analysis using HOMER was also performed leading to identification of ELK1 and ELK4 as candidate upstream transcriptional regulators of differentially upregulated genes in our RNAseq dataset (data not shown). Phosphorylation at key functional sites of ELK-1, e.g., primarily Ser383—sits in the C-terminal transcriptional activation domain—is a direct ERK1/2 substrate^14,19^. Although p38 and JNK can also phosphorylate these sites, experimental evidence indicates that relative to WT MYXV, Δ*M062R* suppresses p38 activation^51^ and does not induce JNK signaling (data not shown). Collectively, these findings support the conclusion that ERK1/2 is the primary upstream activator of ELK-1 during Δ*M062R* infection. The enrichment of growth-related gene transcription, together with ELK1 phosphorylation, provides a mechanistic link between ERK1/2 activation and the transcriptional program induced by Δ*M062R*.

The concurrent activation of ERK1/2 signaling and SAMD9 by Δ*M062R* in the monocytes/macrophages leads to an unusual immunological state that warrants further investigation. In this study, we examined the transcriptional profile after the abortive Δ*M062R* infection and observed a pro-survival and growth promoting effect in these cells. Collectively, our findings suggest that SAMD9 activation does not simply suppress translation indiscriminately. Rather, SAMD9 generates a selective translational environment that preferentially restricts poxviral protein synthesis during Δ*M062R* infection, while concurrent ERK1/2 activation preserves host-cell viability and sustains antiviral immunity. Consequently, these result in a primed antiviral state characterized by heightened responsiveness to subsequent danger signals. These observations expand the current understanding of SAMD9 biology and identify a previously unrecognized interaction between translation control and ERK1/2 pathway in the regulation of innate immunity. As we previously reported, Δ*M062R* treatment in immunosuppressive tumor environment promotes a pro-inflammatory responses in myeloid cells and M2 macrophages^52^. Remodeling of myeloid cell responses subsequently promotes T cell activation leading to effective targeting of tumor cells^52^. In preclinical models, Δ*M062R* has demonstrated activities both as a mono-immunotherapy and in combination with other treatment modalities, such as cisplatin^53^ and DC vaccine^52^, to broaden the therapeutic window and improve therapeutic efficacy for cancer type such as ovarian cancer. Future studies to dissect how Δ*M062R*-induced ERK1/2 signaling compensates translation inhibition by activated SAMD9 pathway leads to the establishment of the potentiated immune state are needed for a better therapeutic utility of this viral agent.

## Data and code availability

This paper does not report original code.

RNA-seq data have been deposited at the NCBI Gene Expression Omnibus (GEO) and will be released upon publication of the manuscript and are publicly available as of the date of publication.

## ACKNOWLEDGMENTS

The study was supported by NIH K22AI099184 and R01AI139106 to JL^#^ (corresponding author), a fund from the River Valley Ovarian Cancer Coalition, a start-up by UAMS Department of Microbiology and Immunology, a UAMS Barton pilot award, and a UAMS ABI-VCRI Collaborative award to JL^#^. This work also received support in part from the Center for Microbial Pathogenesis and Host Inflammatory Responses grant P10GM103625 through the NIH National Institute of General Medical Sciences (NIGMS) Centers of Biomedical Research Excellence (COBRE) and the Winthrop P. Rockefeller Cancer Institute at UAMS to JL^#^. We thank Drs. Martin Cannon and Gaby Fuchs for helpful input, thank Sarah Blair, Bernice Nounamo, Shana Chancellor Owen, and Richard Connor for their technical support of this study. We also thank Drs. Siyuan Ding and Gaopeng Hou for reagents, and Yanling Yang for helpful input.

## AUTHOR CONTRIBUTIONS

Conceptualization, J.L.* (*corresponding author); methodology, J.L.*, E.A.P., and M.L.; Investigation, J.L.*, T.R., E.A.P., L.M., and J.L.; writing – original draft, J.L*.; writing – review & editing, J.L*., T.R., L.M., and J.L.; funding acquisition, J.L*.; resources, J.L*.; supervision, J.L.*.

## SUPPLEMENTAL INFORMATION

Supplemental Figures S1-S2

Supplemental Tables 1-2

---

**Supplemental Figure S1:**
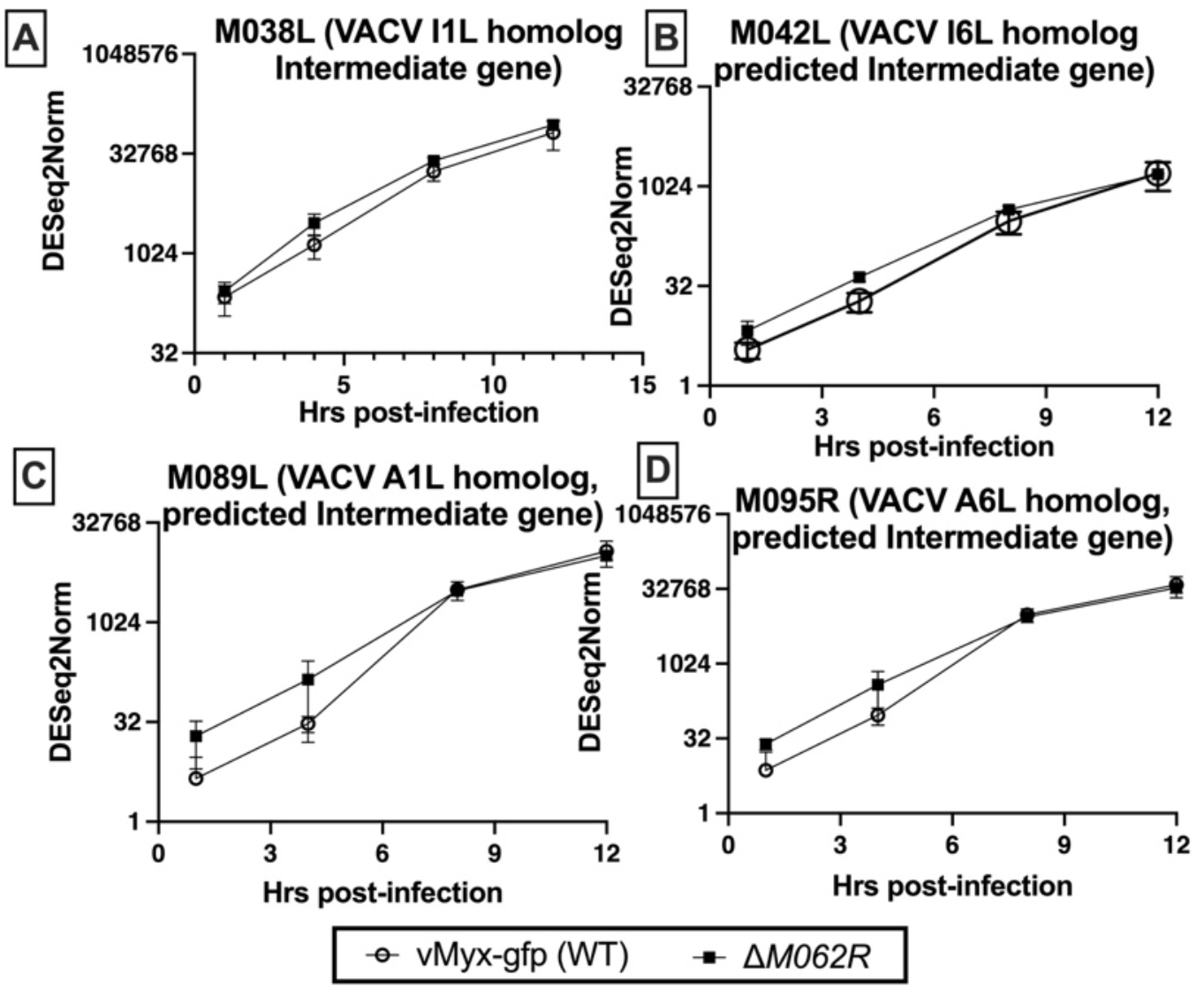
Intermediate gene transcription is not affected by the loss of *M062R* gene during infection. RNAseq was performed using cells that were either infected with WT or Δ*M062R* MYXV at an MOI of 5 for a time course at given timepoints (1, 4, 8, and 12 hrs. p.i.). Three independent replicates were performed and shown is the summary of triplicate data points for each time point and each virus. Known MYXV intermediate genes, (A). *M038L*, (B). *M042L*, (C). *M089L*, and (D). *M095R* were examined for their transcription levels throughout the time course using normalized detection quantification (the median of ratios method in DESeQ2).^54,55,56^

**Supplemental Figure S2:**
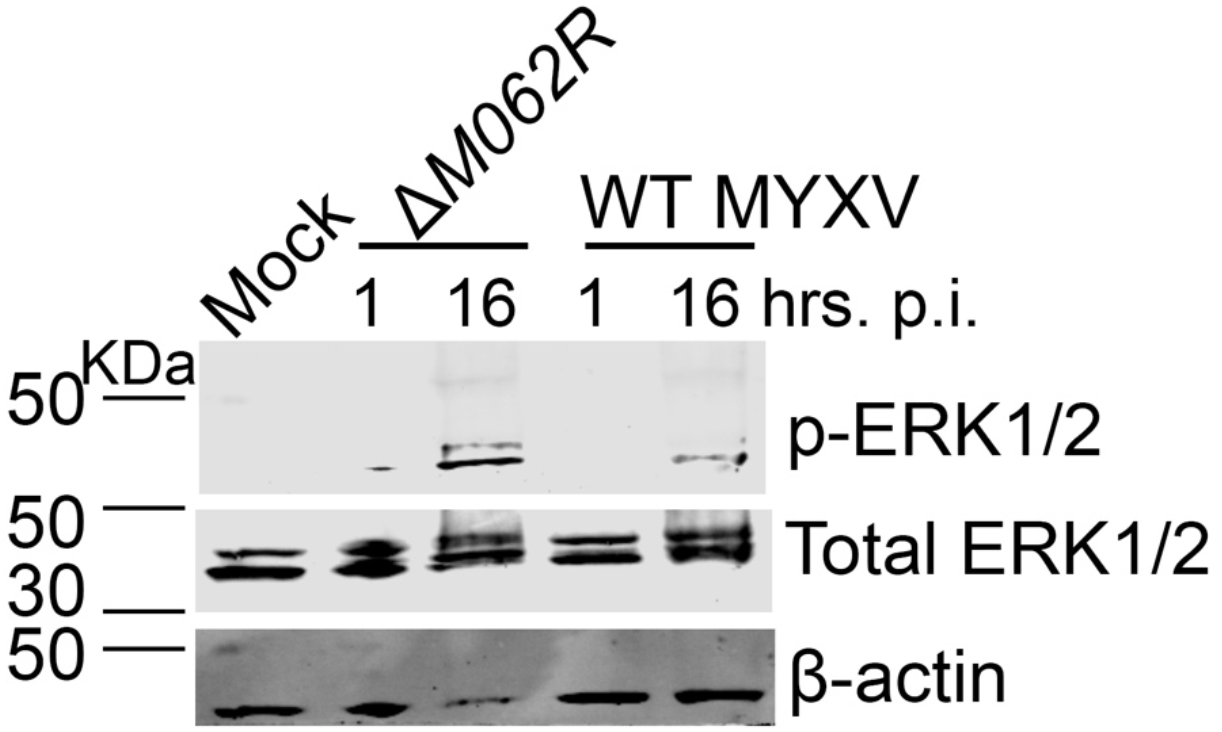
In HeLa cells, D*M062R* infection activates ERK1/2 pathway. HeLa cells were mock treated, infected with Δ*M062R* or WT MYXV at an MOI of 5. At the given time points, 1 and 16 hrs. p.i., cell lysates were harvested for immunoblotting to probe for phosphorylated ERK1/2, total ERK1/2, and β-actin as the loading control.

**Supplemental Table 1.**
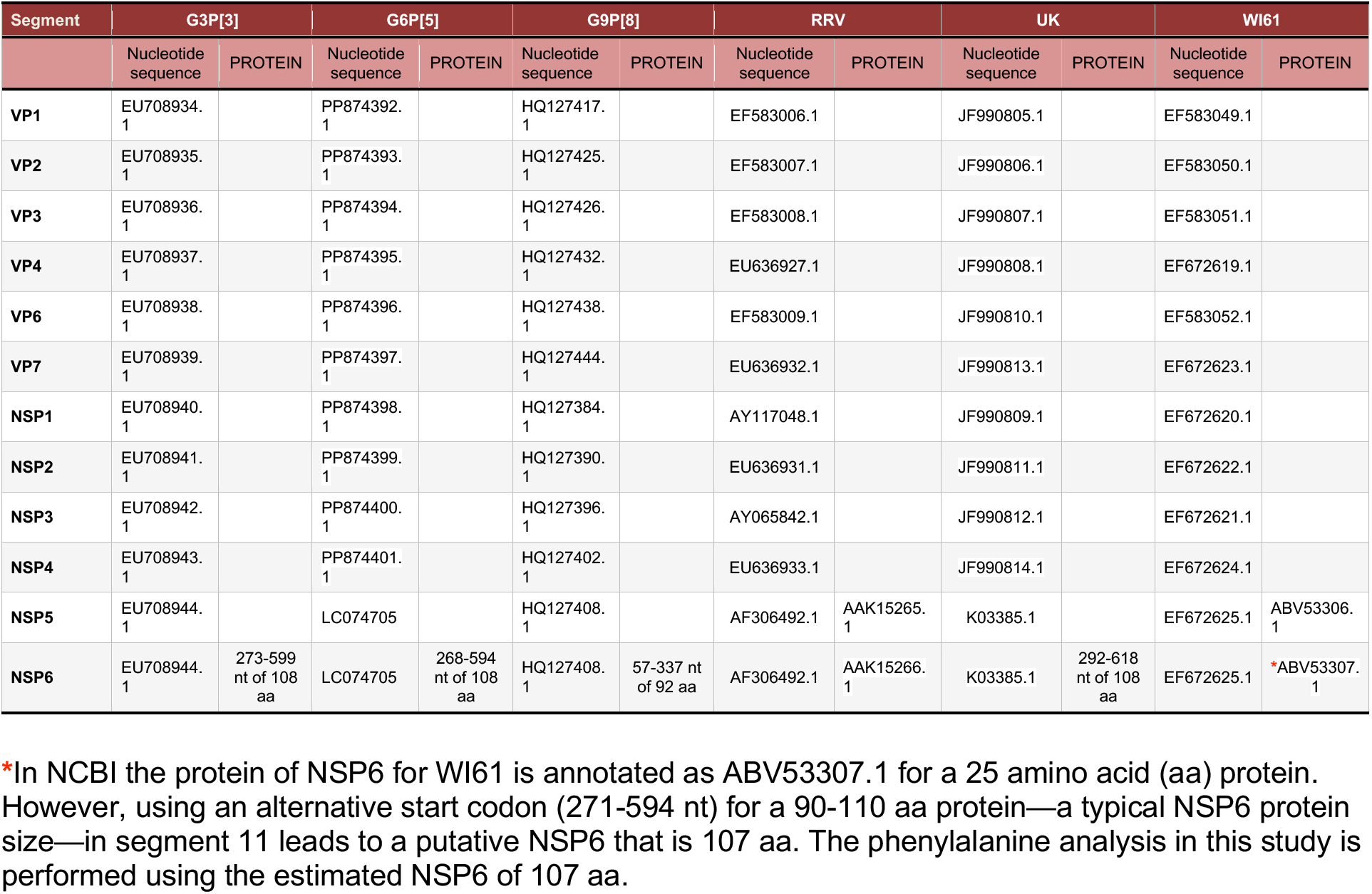
GenBank accession numbers for genome segment nucleotide sequences and protein sequences for rotavirus strains.

| **Segment** | **G3P[3]** |  | **G6P[5]** |  | **G9P[8]** |  | **RRV** |  | **UK** |  | **WI61** |  |
|---|---|---|---|---|---|---|---|---|---|---|---|---|
|  | Nucleotide sequence | PROTEIN | Nucleotide sequence | PROTEIN | Nucleotide sequence | PROTEIN | Nucleotide sequence | PROTEIN | Nucleotide sequence | PROTEIN | Nucleotide sequence | PROTEIN |
| **VP1** | EU708934.1 |  | PP874392.1 |  | HQ127417.1 |  | EF583006.1 |  | JF990805.1 |  | EF583049.1 |  |
| **VP2** | EU708935.1 |  | PP874393.1 |  | HQ127425.1 |  | EF583007.1 |  | JF990806.1 |  | EF583050.1 |  |
| **VP3** | EU708936.1 |  | PP874394.1 |  | HQ127426.1 |  | EF583008.1 |  | JF990807.1 |  | EF583051.1 |  |
| **VP4** | EU708937.1 |  | PP874395.1 |  | HQ127432.1 |  | EU636927.1 |  | JF990808.1 |  | EF672619.1 |  |
| **VP6** | EU708938.1 |  | PP874396.1 |  | HQ127438.1 |  | EF583009.1 |  | JF990810.1 |  | EF583052.1 |  |
| **VP7** | EU708939.1 |  | PP874397.1 |  | HQ127444.1 |  | EU636932.1 |  | JF990813.1 |  | EF672623.1 |  |
| **NSP1** | EU708940.1 |  | PP874398.1 |  | HQ127384.1 |  | AY117048.1 |  | JF990809.1 |  | EF672620.1 |  |
| **NSP2** | EU708941.1 |  | PP874399.1 |  | HQ127390.1 |  | EU636931.1 |  | JF990811.1 |  | EF672622.1 |  |
| **NSP3** | EU708942.1 |  | PP874400.1 |  | HQ127396.1 |  | AY065842.1 |  | JF990812.1 |  | EF672621.1 |  |
| **NSP4** | EU708943.1 |  | PP874401.1 |  | HQ127402.1 |  | EU636933.1 |  | JF990814.1 |  | EF672624.1 |  |
| **NSP5** | EU708944.1 |  | LC074705 |  | HQ127408.1 |  | AF306492.1 | AAK15265.1 | K03385.1 |  | EF672625.1 | ABV53306.1 |
| **NSP6** | EU708944.1 | 273-599 nt of 108 aa | LC074705 | 268-594 nt of 108 aa | HQ127408.1 | 57-337 nt of 92 aa | AF306492.1 | AAK15266.1 | K03385.1 | 292-618 nt of 108 aa | EF672625.1 | *ABV53307.1 |

\*In NCBI the protein of NSP6 for WI61 is annotated as ABV53307.1 for a 25 amino acid (aa) protein. However, using an alternative start codon (271-594 nt) for a 90-110 aa protein—a typical NSP6 protein size—in segment 11 leads to a putative NSP6 that is 107 aa. The phenylalanine analysis in this study is performed using the estimated NSP6 of 107 aa.

**Supplemental Table 2.** Mammalian Orthoreovirus T1L.

| **Mammalian orthoreovirus T1L** |  |  |
|---|---|---|
| **Segment** | **Gene** | **Protein** |
| L1 (rdrp) | M24734.1 | AAA47234.1 |
| L2 (lmbd2) | AF378003.1 | AAK57507.1 |
| L3 (lmbd1) | AF129820.1 | AAD42304.1 |
| M1 (mu2) | X59945.1 | CAA42570.1 |
| M2 (mu1) | AF490617.1 | AAM10735.1 |
| M3 (mu-NS) | AF174382.1 | AAF13169.1 |
| S1 (sigm1) | M35963.1 | AAA47242.1 |
| S1 (sig-1s) | M35963.1 | AAA47243.1 |
| S2 (sig-m2) | L19774.1 | AAA47239.1 |
| S3 (sig-NS) | M14325.1 | AAA47273.1 |
| S4 (sigm3) | M13139.1 | AAA47272.1 |

## KEY RESOURCES TABLE

| REAGENT OR RESOURCE | SOURCE | IDENTIFIER |
|---|---|---|
| Antibodies |  |  |
| β-actin | Sigma Aldrich | A1978 |
| SERP1 | Biomatik | <sup>51</sup> |
| M038 | Biomatik | <sup>51</sup> |
| MDA5 (D6A4) | Cell Signaling Technology | 5321P |
| RIG-I (D14GG) | Cell Signaling Technology | 3743P |
| STAT6 | Cell Signaling Technology | 9362S |
| STAT2 | EMD Millipore | 06-502 |
| OAS2 | Santa Cruz Biotechnology | sc271117 |
| cGAS (D1D3G) | Cell Signaling Technology | 15102S |
| TRIM25 | BD Bioscience | 610570 |
| TRIM21 (D101D) | Cell Signaling Technology | 92043S |
| PKR (B10) | Santa Cruz Biotechnology | sc6282 |
| STING (D2P2F) | Cell Signaling Technology | 13647S |
| IL-1β | R&D | AF-401-NA |
| M062 | Biomatik | <sup>10</sup> |
| Phospho-p44/42 MARK (Erk1/2) (Thr202/Tyr204) (D13.14.4E) | Cell Signaling Technology | 4370S |
| P44/42 MAPK (Erk1/2) (137F5) | Cell Signaling Technology | 4695S |
| Phospho-MEK1/2 (41G9) | Cell Signaling Technology | 9154S |
| MEK1/2 | Cell Signaling Technology | 8727S |
| Phospho-p90RSK (Ser380) (D3H11) | Cell Signaling Technology | 11989S |
| RSK (32D7) | Cell Signaling Technology | 9355T |
| Phospho-eIF4E (Ser209) | Cell Signaling Technology | 9741S |
| eIF4E | Cell Signaling Technology | 9742S |
| Phospho-Mnk1 (Thr197/202) | Cell Signaling Technology | 2111S |
| Mnk1 (C4C1) | Cell Signaling Technology | 2195S |
| Phospho-ELK-1 (Ser383) (2B1) | Cell Signaling Technology | 9186S |
| Elk-1 (F6D8T) | Cell Signaling Technology | 51398S |
| Histone H3 (D1H2) | Cell Signaling Technology | 4499S |
| SAMD9 | Sigma Aldrich | HPA021319 |
| Virus strains |  |  |
| WT MYXV | Dr. G. McFadden, Arizona State University | <sup>8</sup> |
| ΔM062R MYXV | Dr. J. Liu, University of Arkansas for Medical Sciences | <sup>8</sup> |
| Reovirus TD3 | Dr. K. Boehme, University of Arkansas for Medical Sciences | <sup>22</sup> |
| WT VACV | Dr. G. McFadden lab, Arizona State University | <sup>9</sup> |
| VACV-Δ*C7L*/Δ*K1L*-DKO | Dr. J. Liu, University of Arkansas for Medical Sciences | <sup>9</sup> |
| Chemicals, peptides, and recombinant proteins |  |  |
| Streptavidin-HRP | Thermoscientific | 21130 |
| L-Azidohomoalanine (AHA) | Invitrogen | 10102 |
| Benzonase | Millipore Sigma | 70664-3 |
| Biotin Alkyne | Invitrogen | B10185 |
| Phorbol 12-Myristate 13-Acetate | Fisher Scientific | BP685-1 |
| ViaFect | Promega | E4982 |
| Sodium orthovanadate | Santa Cruz Biotechnology | 13721-39-6 |
| Phenylmethylsulfonyl Fluoride (PMSF) | Calbiochem | 52332 |
| β-Glycerophosphate disodium salt hydrate | Millipore-Sigma Aldrich | G5422 |
| Cytosine β-D-arabinofuranoside hydrochloride (Ara-C) | Sigma Aldrich | C6645 |
| Passive Lysis 5X Buffer | Promega | 1910 |
| 2-MERCAPTOETHANOL | MP biomedicals | 60-24-2 |
| Critical commercial assays |  |  |
| Invitrogen™ Inflammation 20-Plex Human ProcartaPlex™ Panel (IL-1β ELISA) | eBioscience/Invitrogen | EXP20012185901 |
| NE-PR Nuclear and Cytoplasmic extraction reagent | Thermofisher Scientific | 78835 |
| Luna Universal qPCR Master Mix | NEB | M3003 |
| TruSeq Stranded Total RNA library prep kit with unique-dual indexing | Illumina | 20020598 & 2002371 |
| cOmplete ULTRA EDTA free Protease inhibitor | ROCHE | 05892791001 |
| CliCK-IT Reaction Buffer | Invitrogen | 10276 |
| Quick DNA/RNA MiniPrep Plus kit | Zymo | D7003 |
| DNeasy Blood & Tissue Kit | QiaGen | 69506 |
| Lipofectamine RNAiMAX | Invitrogen | 13788-075 |
| Methionine and Cysteine free medium | Gibco | 21013024 |
| Deposited data |  |  |
| Experimental models: Cell lines |  |  |
| BSC-40 | ATCC | CRL-2761 |
| HeLa | ATCC | CCL-2 |
| HEK293T | ATCC | CRL-3216 |
| THP-1 | ATCC | TIB-202 |
| THP1-Lucia | Dr. F. Zhu Florida State University | <sup>21</sup> |
| Human CD14<sup>+</sup> monocytes | Lonza Bioscience | 2W-401A |
| Human normal dermal fibroblasts (NHDF) | Lonza Bioscience | CC-2509 |
| THP1-Dual<sup>TM</sup> Cells (NF-κB-SEAP and IRF-Lucia reporter monocytes) | InvivoGen | thpd-nfis |
| THP1-control | Dr. S. Ding, Washington University | <sup>6</sup> |
| THP1-SAMD9-KO | Dr. S. Ding, Washington University | <sup>6</sup> |
| Oligonucleotides |  |  |
| 5’ppp-dsRNA (RIG-I activator) | InvivoGen | tlrl-3prna |
| Software and algorithms |  |  |
| GraphPad Prism | GraphPad Software | Version 10.1.1 |

## METHOD DETAILS

### Phenylalanine bias and phenylalanine codon bias tests

Code was scripted to calculate phenylalanine usage in each gene per genome or cluster of human genes. Relative Synonymous Codon Usage (RSCU) is used to determine whether viral genomes or a cluster of host genes present codon usage bias. RSCU is calculated as the ratio of the observed frequency of a specific code to the expected frequency, which is 1 meaning all synonymous codons for phenylalanine are used with equal probability. When RSCU=1.0, the codon is used equally; RSCU>1.0 suggests the codon is preferred, and RSCU<1.0 indicates the codon is used less frequently than expected with a value under 0.6 classified as strongly underrepresented^57^.

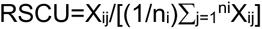

### Cell culture, viruses, Infection, and Titration

Mammalian cells used in this study include BSC-40 (ATCC CRL-2761) (in Dulbecco minimal essential medium, DMEM, Lonza), HeLa (ATCC CCL-2 cultured in DMEM, Lonza), HEK293T (ATCC CRL-3216, DMEM), THP1 (ATCC TIB-202), THP1-dual Lucia (InvivoGen), and THP1-Lucia (kindly provided by F. Zhu Florida State University)^21^ (in RPMI1640, Lonza). Primary normal human CD14^+^ monocytes (Lonza, Walkersville, MD) (RPMI1640) and primary normal dermal fibroblasts NHDF (Lonza, Walkersville, MD) (5% FBS-DMEM) were also used in this study. The complete growth medium (e.g., DMEM Lonza/BioWhittaker Catalog no 12-604Q, or RMPI1640) was supplemented with 10% FBS (Atlanta Biologicals, Minneapolis, MN) unless specified otherwise, 2 mM glutamine (Corning Cellgro, Millipore Sigma, St. Louis, MO), and 100 μg per ml of Pen/Strep (Corning Cellgro, Millipore Sigma, St. Louis, MO); for RPMI1640 complete culture medium, in addition to FBS, glutamine, and Pen/Strep, 2-mercaptoethanol (MP biomedicals, Solon, OH) was supplemented to a final concentration of 0.05 mM.

The myxoma virus (MYXV) used in the study is from Lausanne strain background (GenBank Accession AF170726.2).^8^ The *M062R*-null MYXV (Δ*M062R*) targeted deletion mutant and wildtype virus (WT-GFP) were described previously.^8^ Vaccinia virus used in this study is from WR strain background (GenBank Accession AY243312.1). Both WT VACV and VACV-Δ*C7L*/Δ*K1L* (double knockout virus, DKO) have been described previously^9^. Poxvirus stocks were amplified on BSC-40 cells and purified with sucrose step gradient through ultracentrifugation as previously described.^58^ Infection is performed by incubating virus at appropriate MOI in minimal volume of inoculum media (1% FBS diluted in PBS) on ice or at 37C under frequent rocking for 1 hour before washing cells with PBS to rid unbound viral particle before replacement with normal growth medium. Reovirus TD3 was provided by K. W. Boehme (University of Arkansas for Medical Sciences) and described previously.^22^ Poxvirus titration is as previously described using BSC-40 cells^8,9^.

### Western blot and Multi-plex Cytokine Array

THP-1 cells, including parental THP1, control, and SAMD9-null THP1 cells, are first differentiated with PMA (FisherScientific, Cat# BP685-1) at 50 ng/ml for up to 48 hours before mock treated, infection with WT, Δ*M062R* at a MOI of 5. To inhibit post-replicative gene expression, Ara-C of 100 and 200 μM is introduced 1 hour before infection till the time of harvesting. For experiment examining VACV effect, WT VACV and VACV-Δ*C7L*/Δ*K1L* are used to infect cells at a MOI of 5. NHDF and HeLa cells are also used for immunoblotting after infection with corresponding viruses as specified in each experiment. Cells were harvested at given time points with the NP-40 lysis buffer [1% NP-40, 50mM Tris-Cl pH 7.4, 150mM NaCl containing the protease inhibitor (cOmplete ULTRA EDTA-free Protease inhibitor, Cat# 6538282001, MilliporeSigma), 2 mM sodium orthovandate (Santa Cruz Biotechnology, Inc, Cat#13721-39-6), 1 mM PMSF (Cat# 52332, MilliporeSigma), and 25mM β-Glycerophosphate (SigmaAldrich, Cat# G9422)] or RIPA lysis buffer (for time course experiment probing ELK-1 signaling). Cell lysates were incubated on ice for 30 minutes before centrifugation at 125,000 g for 10 min and 50 μg of total protein for each sample was separated on SDS-PAGE [Mini-PROTEAN TGX Gels (Biorad, Cat# 456-1095)]. Proteins were transferred to 0.2 um Nitrocellulose membrane (Cat# 1620112) for antibody probing. To examine effector signaling of both cytoplasmic (e.g., MER1/2, p90RSK, eIF4E, and MNK1, etc.) and nuclear (ELK-1) proteins at the same time, after corresponding viral infection at a MOI of 5 at 18 hrs. p.i., cells are harvested with NE-PER Nuclear and Cytoplasmic Extraction Reagents (Thermo Fisher Scientific, Cat# 78835) containing cOmplete ULTRA EDTA-free Protease inhibitor, Cat# 6538282001) along with 2mM sodium orthovanadate (Santa Cruz, Cat# 13721-39-6), 1mM PMSF (Millipore Sigma, Cat# 52332) and 25mM β-Glycerophosphate (sigma Aldrich, Cat# G9422) according to manufacturer instructions. Protein concentration for samples extracted with NE-PER Nuclear and Cytoplasmic Extraction kit is measured using BCA reagents; 30 μg of total protein from the cytoplasmic fraction and 15 μg from the nuclear fraction are used for immunoblotting. For experiments examining ERK1/2 effect using inhibitor of ERK signaling, U0126, treatment of U0126 at a dose of 10 μM is added after infection at given time point for 3 hours before harvesting for separation of cytoplasmic and nuclear protein using NE-PER kit. Quantification of immunoblotting signals is performed using Image Studio^TM^ version 6.2.0.116 (LI-COR Biotech) or Image Lab Software version 6.1.0 build 7 ChemiDoc MP (Bio-Rad). Phosphorylated protein signals from each group is first normalized to corresponding total protein from the same group; to obtain standardized signaling, the resulting ratio from treatment group is then normalized to uninfected cells. For each blot, independent measurement in triplicate are averaged for statistical comparison and for calculation of standard deviation (STDEV).

For detection of IL1α, IL1β, and IL-8 in the supernatant, the Human Inflammation Panel (Invitrogen™ Inflammation 20-Plex Human ProcartaPlex™ Panel) (Catalog # EXP20012185901, eBioscience/Invitrogen) was used following manufacturer standard protocol with the BioRad Bio-Plex 200 system (Bio-Rad, Hercules, CA).

### Pulse-labeling to detect newly synthesized protein

Differentiated THP1 cells were infected with wildtype or Δ*M062R* MYXV at an MOI of 5 to 16 hrs before culturing in medium free of methionine and cystine (Gibco, Cat# 21013024) for 1-2 hr. Pulse-labeling was performed with 25 µM AHA (L-Azidohomoalanine) (Invitrogen, Cat# 10102) for 2 hrs before cell lysate harvesting in 500 µl of RIPA lysis buffer (1% NP-40, Tris pH 7.4, NaCl 150mM, 0.5% Sodium deoxycholate, 0.1% Sodium dodecyl sulfate, and the protease inhibitor). Cell lysates were treated with 250U of Benzonase (Novagen, Cat# 70664-3) and incubate for 30 minutes before centrifugation at 12,500xg 4°C for 10 min, 200 µg total proteins were subjected to Click-iT chemistry reaction using 100 µL of Click-iT Reaction Buffer (Invitrogen, Cat# 10276) and 40 µM of Biotin Alkyne (Invitrogen, Cat# B10185) following manufacturer’s instructions. After precipitation (Methanol and chloroform) 20 µg of resolubilized protein (1% SDS in 50 mM Tris-HCl pH8.0) from each sample was analysed in Western blot probed with Streptavidin-HRP (Thermo scientific, Cat# 21130).

### Realtime PCR to examine viral DNA replication

Cells were infected with either WT MYXV or Δ*M062R* at a MOI of 5. At the given time points (4, 8, 12, 24 hrs. p.i.) DNA extraction was performed using DNeasy Blood & Tissue Kit (Qiagen, Cat. No. 69506) followed by realtime PCR (Luna Universal qPCR Master Mix, Cat. No. M3003, NEB Inc) following manufacturer standard protocol. Realtime PCR primers recognizing viral *M071L* and host 18s were used for quantification of viral DNA levels as previously described.^8^

### RNAseq and data processing

THP-1 cells were differentiated using PMA at 50 ng/mL for up to 48 hours before mock treated, transfection with interferon stimulated DNA (ISD)^59^ at 2 μg per dish using ViaFect based on manufacturer protocol (Cat. No. E4982, Promega, Madison, WI), or infected at an moi of 5-10 for either WT MYXV or Δ*M062R*. At 1-, 4-, and 8-hours post-transfection of ISD and 1-, 4-, 8-, and 12-hours post-infection cells were harvested and spun down for RNA extraction. RNA extraction was performed as previously described.^10^ Briefly, after Quick DNA/RNA MiniPrep Plus kit (Cat. No. D7003; Zymo, Irvine, CA, USA), quality control of purified RNA was performed using the Qubit RNA BR Assay kit (Cat. No. Q10211; Invitrogen, Waltham, MA, USA) and the Standard Sensitivity RNA Analysis kit on a Fragment Analyzer capillary electrophoresis system (Cat. No. DNF-471-0500; Agilent, Santa Clara, CA, USA).

TruSeq Stranded Total RNA library prep kit with unique-dual indexing (Cat. No. 20020598 & 2002371; Illumina, San Diego, CA, USA) was used for this study with 250 ng of total RNA. Quality control of the libraries was conducted using the Qubit 1X dsDNA HS Assay kit (Cat. No. Q33231; Invitrogen, Waltham, MA, USA) for mass, the High Sensitivity NGS Fragment Analysis kit on a Fragment Analyzer capillary electrophoresis system (Cat. No. DNF-474-0500; Agilent, Santa Clara, CA, USA) for fragment size, and the Universal Library Quantification kit (Cat. No. 07960140001; KAPA, Wilmington, MA, USA) for functional validation. Finally, validated libraries were adjusted to 3 nM followed by pooling, denaturing, and clustering. HiSeq 3000 (Illumina, San Diego, CA, USA) was used for paired end (2X75) sequencing to an average of 40 million reads per sample.

After QC of raw FASTQ files, a dualRNAseq pipeline is used for sample alignment as previously described.^10^ Normalization and differential analysis is performed using using R v4.5.1^60^ and DESeq2 v1.49.5 with an adjusted p value ≤0.05 defined as the significance threshold.^61^ Genes with *log₂FoldChange* > 0 were classified as upregulated, and those with *log₂FoldChange* < 0 as downregulated. The time-course RNA-seq analysis was conducted using maSigPro v1.81.0.^62^ Normalized expression data from DESeq2 were used to identify gene clusters exhibiting significant temporal expression changes. Gene expression comparisons within figures were made by normalizing raw count data using the median of ratios method in DESeQ2.^54,55,56^ The Reactome pathway enrichment plot were generated using the R library Reactome PA v1.44.0 and the *enrichPathway* function.^63^ Only upregulated significant differentially expressed genes were passed to the aforementioned function and the top 10 pathways are displayed.

### Luciferase assay to examine viral intermediate gene expression

HeLa cells were transfected with the plasmid containing the luciferase expression cassette driven by one of following VACV intermediate gene promoters, pA2L,^11^ pI1L,^11^ and VACV intermediate consensus promoter sequence, pINT^64^ (kindly provided by M. Wiebe, University of Nebraska-Lincoln), using ViaFect (Cat# E4982, Promega, Madison, WI) at 0.8 μg/mL with the transfection agent to DNA ratio of 3 μL to 1 μg as previously described.^10^ At 16-24 hours post-transfection, cells were infected with either WT or Δ*M062R* MYXV at a MOI of 5 in the presence or absence of 100-200 μM Ara-C (Sigma-Aldrich, C6645) for 18 hours before harvesting with Passive Lysis Buffer (Cat# 1910, Promega) and measuring luciferase activities using Luciferase Assay System (Cat# E1500, Promega) following manufacturer instructions.

### Growth and viability comparison

The same numbers of THP1 cells are mock treated or infected with corresponding viruses (WT MYXV, Δ*M062R*, WT VACV, and VACV-Δ*C7L*/Δ*K1L*) at a MOI of 1. After wash to get rid of unbound virus, cells are seeded at 2E+05 cells per mL. At given time points, live and dead cells are counted using Trypan blue exclusion assay to estimate live cell growth in triplicate of technical replicate. Shown is a representative growth comparison of 3 biological replicates.

### Luciferase-based assay to assess type 1 interferon induction

THP1 cells with IRF core promoter driven luciferase expression or THP1-dual cells (IRF-luciferase and NF-κB-SEAP, Invivogen, Cat. No. thpd-nfis) were differentiated into macrophages as previously described.^10^ After pretreatment of mock infection, infection with WT MYXV or Δ*M062R* at an MOI of 5 for 9 hours before exposure to reovirus TD3^22^ at an MOI of 50 or 5’ppp-dsRNA (RIG-I activator) (Invivogen, Cat. No. Tlrl-3prna) transfected with Lipofectamin RNAiMAX (Invitrogen, Cat. No.13788-075). At 12 hours after the secondary infection by reovirus or exposure to stimuli such as 5’ppp-dsRNA, supernatant was harvested for luciferase assay (QUANTI-Luc, Invivogen, Cat. No. rep-qlc1) following manufacturer’s standard protocol. The fold induction is shown which is normalized to baseline signal as previously described^10^ and the summary of 3 biological replicates with duplicate of technical replicates.

## QUANTIFICATION AND STATISTICAL ANALYSIS

GraphaPrism 10.1.1 was used for statistical analyses. One-way ANOVA followed by multiple comparisons (i.e, Tucky’s test following the Ordinary one–way ANOVA), two–way ANOVA, or Student t test or multiple paired t test was performed according to experiment setup. Statistical significance is defined as *p<0.05 (**p<0.01, ***p<0.001, and ****p<0.0001).

